# CD164 is an endolysomal host factor for entry of Clade A New World Arenaviruses

**DOI:** 10.64898/2026.04.21.719929

**Authors:** Cassandra E. Thompson, Tomasz Kaszuba, Rita M. Meganck, Marjorie Cornejo Pontelli, Paul W. Rothlauf, Mason A. Roth, Zhuoming Liu, Hongming Ma, John Doench, Michael S. Diamond, Daved H. Fremont, Sean P.J. Whelan

## Abstract

Arenaviruses are divided into Old World (OW) and New World (NW) groups. OW arenaviruses enter cells through a pH-dependent receptor switch from a plasma-membrane factor to an endolysosomal receptor for subsequent membrane fusion, whereas clade B NW arenaviruses use transferrin receptor 1 without a secondary receptor. Using a vesicular stomatitis virus (VSV) chimera expressing the glycoprotein complex (GPC) of the clade A NW arenavirus Pichindé virus, we performed a genome-wide CRISPR loss-of-function screen and identified the endolysosomal sialomucin CD164 as an essential host factor. CD164 knockout cells were resistant to VSV chimeras bearing the GPCs of Pichindé, Paraná, and Flexal viruses, and to authentic Pichindé and Paraná virus, with susceptibility restored by complementation. The requirement mapped to the cysteine-rich domain of CD164, which bound GP1 in a pH-dependent manner through main-chain interactions. These findings define CD164 as an endolysosomal receptor for clade A NW arenaviruses expanding the receptor switching paradigm.

## Introduction

Arenaviruses are globally distributed zoonotic pathogens that persist in rodent reservoirs and can cause severe disease in humans following exposure to contaminated rodent excreta or secretions.^1–3^ Several members of the family *Arenaviridae*, including Lassa virus and the South American hemorrhagic fever viruses, produce life-threatening infections and are classified as Category A priority pathogens due to their epidemic potential, high case-fatality rates, and lack of effective therapeutics and vaccines.^4,5^ Mammalian arenaviruses are divided into Old World (OW) and New World (NW) groups according to their geographic distribution, with NW arenaviruses (NWA) further subdivided into clades A-D (**Fig. 1A**).^6,7^ The clade A NWA are not typically associated with severe human disease, but have been implicated in human infection and seroconversion, highlighting a continued need to understand their biology and potential for emergence.^7,8^

**Figure 1:**
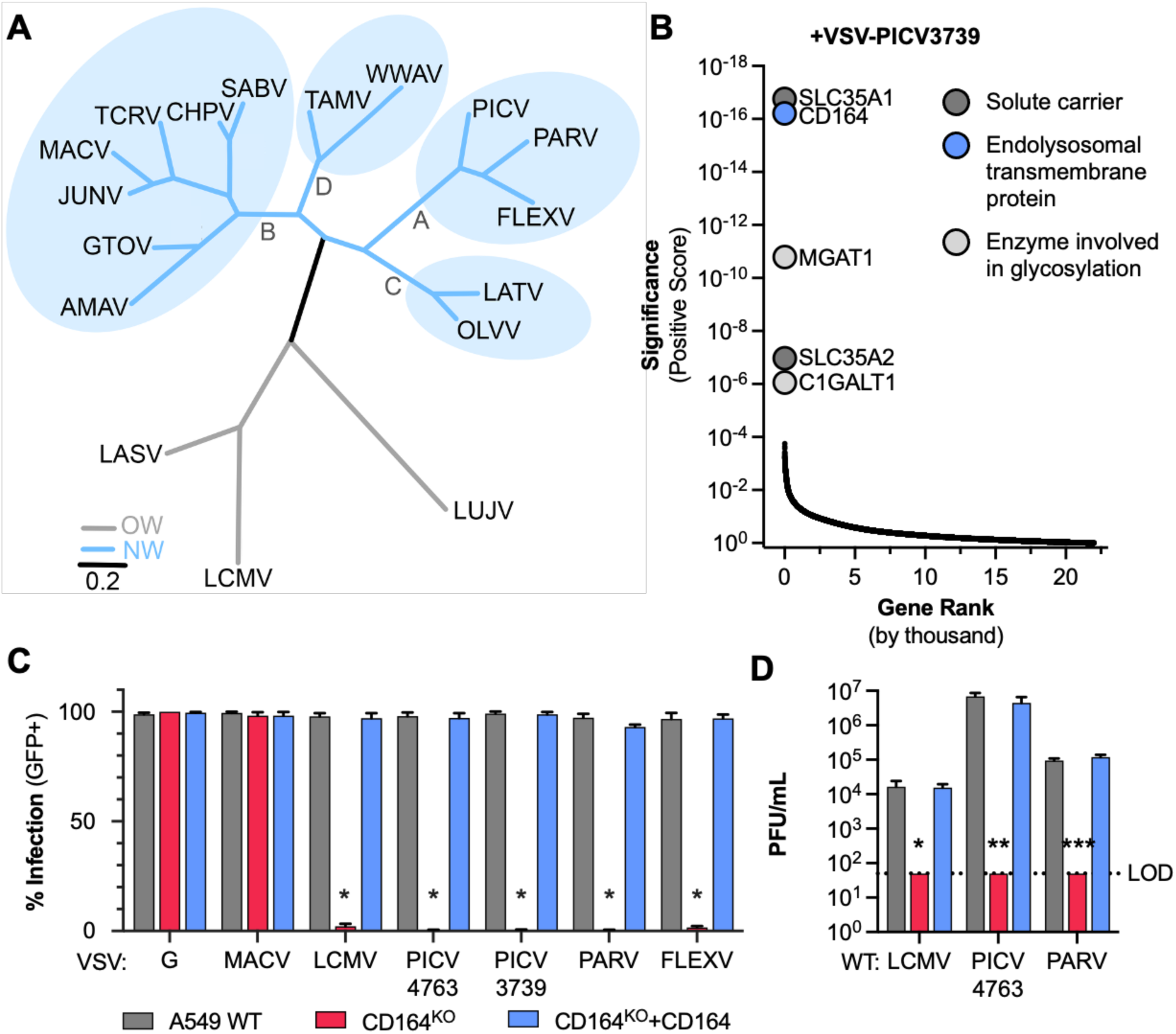
Genetic screens identify host factors required for Clade A New World arenavirus infection. **(A)** Phylogenetic tree of mammarenavirus GPC amino acid sequences. Tree generated via MUSCLE alignment, Gblock curation, PhyML phylogeny, and rendered with TreeDyn using phylogeny.fr. **(B)** Enrichment of guide RNAs from a CRISPR-Cas9 loss-of-function genome-wide screen after the second round of infection with chimeric VSV-eGFP-PICV (CoAN3739). Genes arranged by rank on the x-axis; significance (positive score) on the y-axis. (n=1, experimental replicate). **(C)** WT A549, CD164^KO^, and CD164^KO^+CD164 cells were infected with WT or chimeric VSV-eGFP expressing indicated glycoproteins (MACV, LCMV, PICV CoAN4763, PICV CoAN3739, PARV, FLEXV) at MOI=3. Six hours post-infection, % GFP-positive cells determined by flow cytometry (n=3, independent experiments in triplicate and technical replicates in duplicate). One-Way ANOVA: \**p*<0.05, \*\**p*<0.005, \*\*\**p*<0.0005. **(D)** WT A549, CD164^KO^, and CD164^KO^+CD164 cells were used for plaque assays with WT LCMV (Armstrong), PICV (CoAN4763), and PARV. Dotted line indicates limit of detection (LOD). (n=3, independent experiments in triplicate). One-way ANOVA with Tukey’s multiple comparisons: *\*p<*0.05, *\*\*p<*0.005, *\*\*\*p<*0.0005.

Arenaviruses are enveloped, bi-segmented, negative-sense RNA viruses whose surface is decorated by a glycoprotein complex (GPC), a class I viral fusion protein required for cell entry.^9,10^ GPC is cleaved by the host signal peptidase^11^ and also by Subtilisin kexin isozyme 1 (SKI-1)/site 1 protease (S1P) in the Golgi,^12–14^ to yield a stable signal peptide (SSP; residues 1-58), GP1 (residues 59-272), GP2 (residues 273-508), that form a heterotrimeric spike that mediates attachment and membrane fusion.^15–20^ Entry begins when GP1 engages a host factor at the plasma membrane, with subsequent endocytosis and transport to endolysosomal compartments. Within those compartments, acidic pH triggers conformational rearrangements in the glycoproteins that culminate in GP2-mediated fusion of viral and endolysosomal membranes to release the viral ribonucleoproteins into the cytoplasm.^21^

For Lassa virus (LASV) and lymphocytic choriomeningitis virus (LCMV), two OW arenaviruses (OWA), GP1 binds α-dystroglycan (α-DG) at the plasma membrane,^22–25^ and undergoes an acid pH-dependent switch to their respective endolysosomal receptors LAMP1 for LASV,^26^ and CD164 for LCMV.^27,28^ Lujo virus, another OWA, undergoes a similar pH-dependent switch from neuropilin 2 (NRP2) at the plasma membrane to CD63 in the endolysosome.^29,30^ These pH-dependent receptor switches are essential for productive fusion and are a shared feature of OWA entry. The best characterized entry mechanisms for the NWA are exemplified by the pathogenic clade B viruses, such as Machupo virus (MACV), which binds transferrin receptor 1 (TfR1) at the cell surface.^31–35^ Although fusion of NWA also occurs from late-endosomes or lysosomes, host factors required for that process have not been identified. Additional host factors have also been implicated in NWA attachment to cells including phosphatidylserine receptors and C-type lectins.^36,37^ Voltage-gated calcium channels (VGCCs) promote Junín virus (JUNV) and Tacaribe virus (TACV) infection by enhancing viral attachment independent of TfR1.^38,39^

Here we employed chimeric vesicular stomatitis viruses (VSVs) bearing clade A NWA GPCs to perform unbiased CRISPR-Cas9 loss-of-function screens for host factors required for infection. This approach identified CD164 as an essential factor for infection by multiple clade A NWA. Like LCMV, we demonstrate that the GP1 of several clade A NWA bind directly to the cysteine-rich domain (CRD) of CD164 in an acid pH dependent manner, leading to virus-cell fusion. Our results define a unique endolysosomal receptor for NWA and reveal unifying principles of receptor usage and fusion across *Arenaviridae*.

## Results

### Genetic screens identify host factors for clade A NWA infection

To identify host determinants of clade A NWA entry into cells, we generated chimeric vesicular stomatitis viruses (VSV) in which the native glycoprotein (G) was replaced by the glycoprotein complex (GPC) of Pichindé virus (PICV), Paraná virus (PARV), or Flexal virus (FLEXV) **(Fig 1A, S1A)**. All of the VSV chimeras encoded enhanced green fluorescent protein (eGFP) to enable a direct visual readout of infection **(Fig S1A)**. Each of the VSV-GPC chimeras grew to reduced titer and exhibited smaller foci of infection than wild-type VSV **(Fig S1B)**, consistent with attenuation typical for replacement of G by the arenavirus GPC.^40–42^ Incorporation of the respective GPC into viral particles was confirmed biochemically **(Fig S1C)**, and by electron microscopy **(Fig S1D)**.

We performed a genome-wide CRISPR-Cas9 loss-of-function screen in Vero CCL-81 cells using the VSV-PICV CoAN3739 GPC chimera (VSV-PICV3739). After two rounds of selection by infection, deep sequencing of enriched guides identified multiple candidate host factors **(Fig 1B, Table S1)**. Among the top hits were genes involved in glycosylation and sialyation (*SLC35A1*, *SLC35A2*, *MGAT1*, *C1GALT1*), and *CD164*, a highly glycosylated sialomucin previously implicated as a receptor for LCMV.^27,28,43,44^ We carried out three additional CRISPR-Cas9 loss-of-function screens in a haploid human cell line (HAP1) using VSV-PICV3739, VSV-PARV and VSV-FLEXV and a lentivirus library of 4,630 single guide RNAs (sgRNAs) targeting 1,146 human genes that encode predicted plasma-membrane proteins. After one round of infection at an MOI of 3, selected cells were non-permissive to further infection and enriched sgRNAs were identified by deep sequencing (**Fig S2A**, **Table S2**). For each virus, CD164 was ranked as the top hit, and given its known role in LCMV infection, we prioritized this hit for further characterization.

To investigate the requirement for CD164, we compared infection in A549 cells, CD164 knockout (CD164^KO^) cells, and CD164^KO^ cells complemented by overexpression of human CD164 (CD164^KO^+CD164).^27^ Infection by wild type VSV, and VSV-MACV (clade B NWA) were unaffected by loss of CD164 **(Fig 1C)**. By contrast, infection by VSV-LCMV and all clade A NWA including VSV-PICV CoAN4763, VSV-PICV3739, VSV-PARV, and VSV-FLEXV was inhibited by >95% by knockout of CD164 and was fully restored by complementation **(Fig 1C)**. To validate these findings with authentic arenaviruses, we infected A549 cells with wild-type LCMV Armstrong, PICV CoAN4763, and PARV. As with the VSV-chimeras, infection was ablated in CD164^KO^ cells and rescued by CD164 complementation **(Fig 1D)**. Together, these results demonstrate that CD164 is essential for GPC-mediated entry of clade A NWA and that VSV-chimeras can be used to identify host factors required for GPC-mediated infection.

### The cysteine-rich domain of CD164 is required for clade A NWA infection

The ectodomain of CD164 comprises two mucin domains (MD1, MD2) flanking a central cysteine-rich domain (CRD) **(Fig 2A**).^27,44^ Domain mapping in CD164^KO^ Hela cells revealed that deletion of MD1 (ΔMD1) had no effect on infection by clade A NWA or LCMV VSV-chimeras, whereas deletion of either the CRD (ΔCRD) or MD2 (ΔMD2) ablated infection **(Fig 2B)**. To discrimintate between a direct role of MD2 from effects on CRD presentation, we fused the CRD to the luminal stalk of LAMP1 to ensure proper endolysosomal targeting (CRD-LAMP1, **Fig 2A**). This CRD-LAMP1 fusion fully restored infection by all clade A NWA and LCMV chimeras, establishing that the CRD alone is the functional determinant required for GPC mediated entry **(Fig 2B)**.

**Figure 2:**
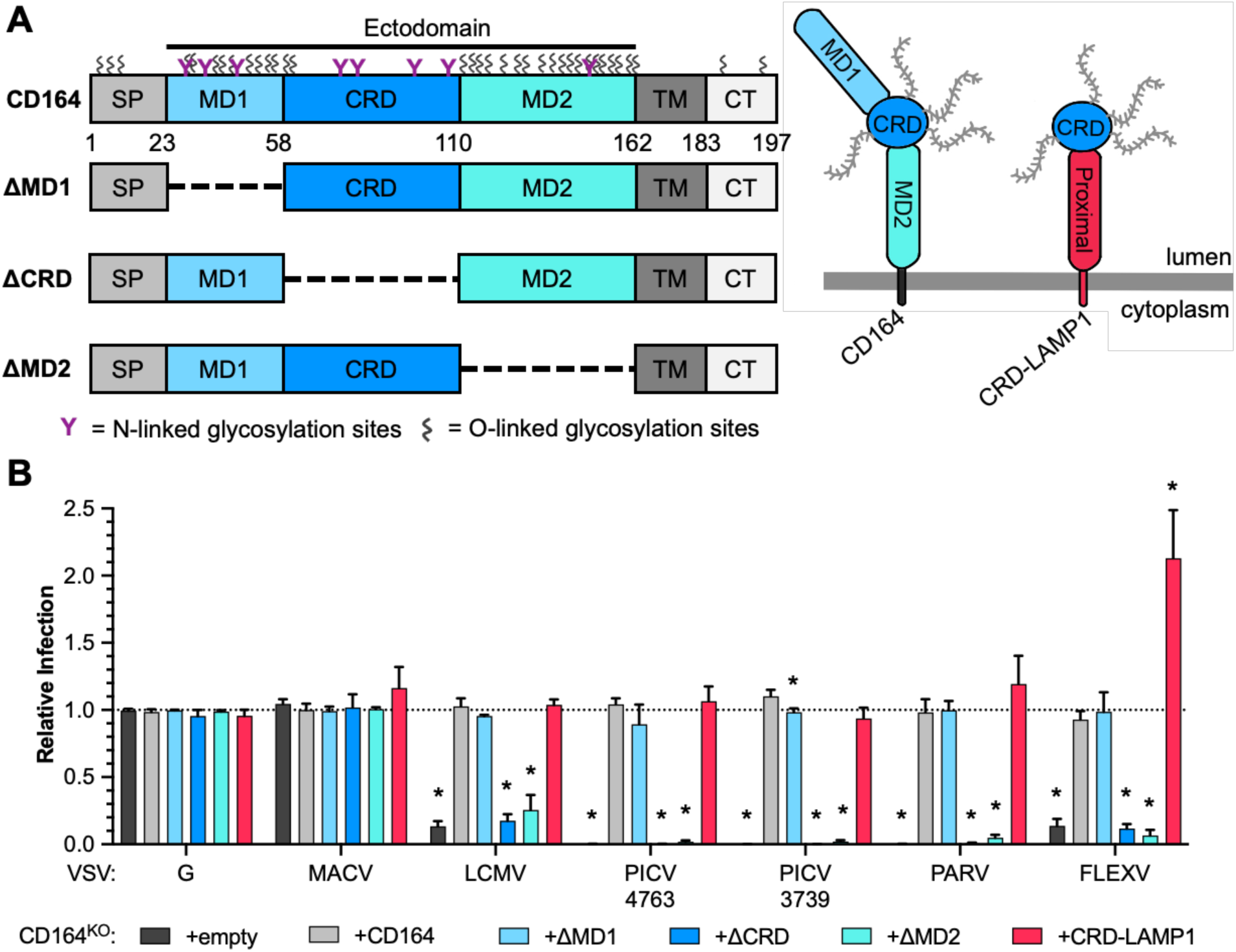
Domain mapping of CD164 requirements for clade A NWA GPC-mediated infection. **(A)** Left: domain architecture of CD164 and deletion mutants; predicted N-linked glycosylation sites (purple), O-linked sites (grey). Right: schematic comparing CD164 and CRD-LAMP1 chimeric protein (CD164 CRD: blue; LAMP1 proximal domain: red). **(B)** CD164^KO^ HeLa cells expressing empty vector, CD164, CD164 deletion mutants (ΔMD1, ΔCRD, ΔMD2), or CRD-LAMP1 were inoculated with chimeric VSV-eGFP expressing indicated glycoproteins (MOI=2). % GFP-positive cells determined 6 hpi by flow cytometry; normalized to WT HeLa cells (Relative Infection) (n=3, independent experiments in triplicate and technical replicates in duplicate). One-Way ANOVA: \**p*<0.05.

### CD164 and acidic pH are required for glycoprotein-mediated fusion

For LASV, LCMV, and LUJV, endolysosomal receptors facilitate the GP2-dependent fusion of virus with the host membrane.^26–29^ We therefore next examined whether CD164 acts at the pH-dependent fusion step for clade A NWA using an acid-bypass assay. PICV GPC initiates plasma membrane fusion at pH 5.4 when viral particles are bound to CD164^KO^ cells complemented with plasma membrane-targeted CD164 (+WT-CD164^PM^) at 4°C and exposed to acidic pH **(Fig 3A)**. By contrast, VSV-chimeras containing the GPC of LCMV, PICV3739, PARV or FLEXV bound to CD164^KO^ cells and exposed to pH 5.2 were defective in infection unless cells were complemented with plasma membrane-targeted CD164 **(Fig 3B)**. As expected, fusion and infection of cells by VSV-MACV was unaffected by the presence or absence of CD164 **(Fig 3B)**. These results demonstrate that CD164 is required for acid-triggered membrane fusion for clade A NWA.

**Figure 3:**
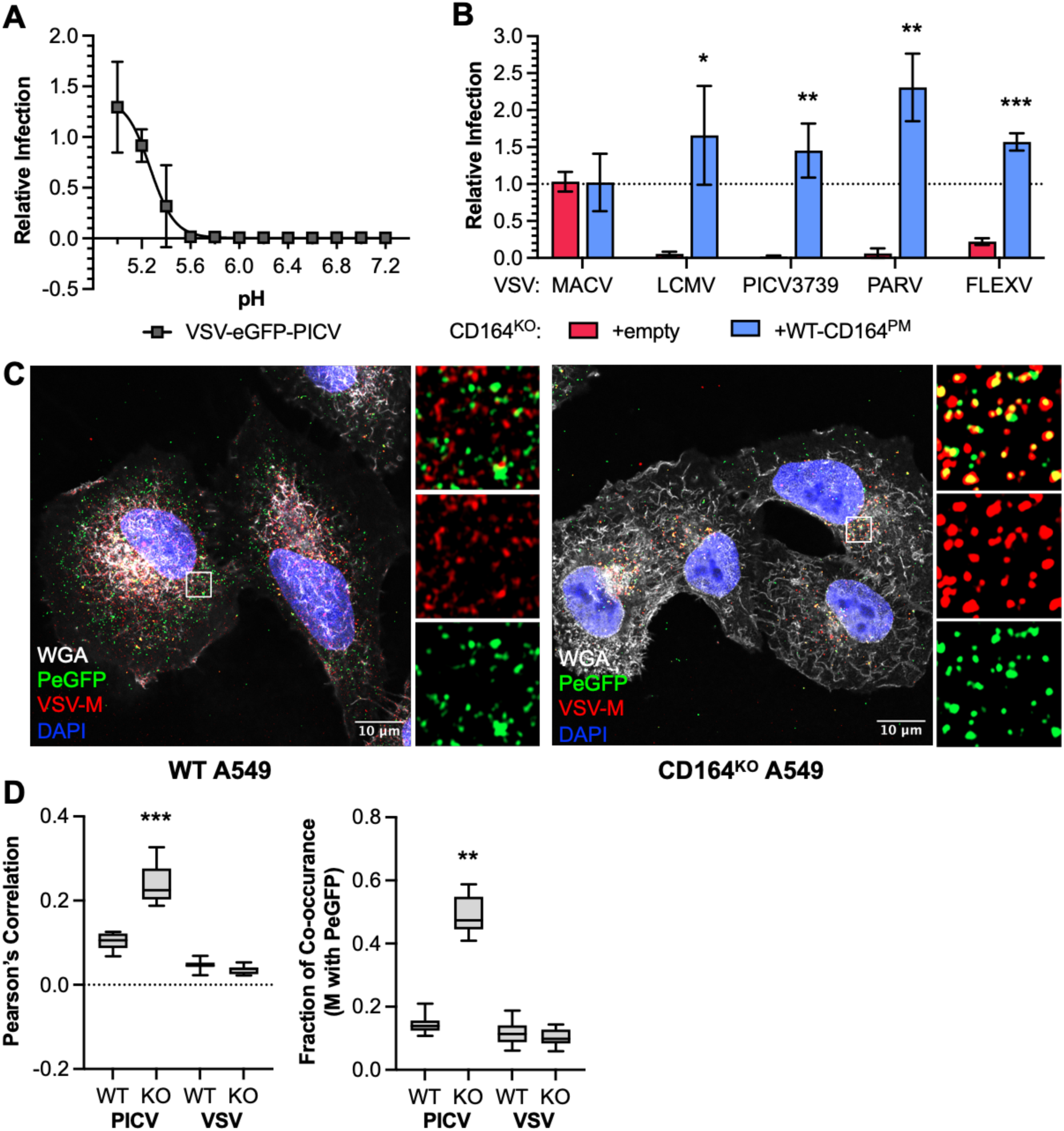
Effect of CD164 on GPC-mediated fusion. **(A)** Acid by-pass assay in CD164^KO^ HeLa cells expressing plasma membrane-targeted CD164 (WT-CD164^PM^). VSV-eGFP-PICV3739 was bound to Bafilomycin A1-pretreated cells (MOI=3, 4°C, 1 hr) then fusion was triggered at indicated pH, media replaced, and infection proceeded for 5 hr in Bafilomycin A1. % GFP-positive cells normalized to WT HeLa at pH 5.0 (n=3, independent experiments in triplicate and technical replicates in duplicate). **(B)** Same assay as in (A) performed with WT, CD164^KO^ cells expressing empty vector, or WT-CD164^PM^ cells, with fusion triggered at pH 5.2. % GFP-positive cells normalized to WT HeLa (dotted grey line) (n=3, independent experiments in triplicate and technical replicates in duplicate). One-Way ANOVA: \**p*<0.05, \*\**p*<0.005, \*\*\**p*<0.0005. **(C)** WT or CD164^KO^ A549 cells pretreated with cycloheximide were inoculated with VSV-P/eGFP-PICV3739 (MOI=400, 3 hr). Cells were fixed, stained with DAPI, WGA, anti-VSV-M, and imaged by Airyscan confocal microscopy. Representative cell shown per condition. VSV-M: red; P/eGFP: green; colocalization: yellow; WGA: white; DAPI; blue. Scale bar, 10 µm. **(D)** Quantification of P/eGFP and VSV-M colocalization from (C) (20 cells per condition). One-Way ANOVA (left) or Kruskal-Wallis ANOVA (right), \**p*<0.05, \*\**p*<0.005, \*\*\**p*<0.0005..

To independently assess whether CD164 was required for membrane fusion, we capitalized on a single-particle content-release assay using VSV-chimeras in which eGFP is fused to an internal structural component of the virus particle – the phosphoprotein (P) – and separately detect the viral matrix protein (M) using a well-characterized monoclonal antibody 23H12.^45–47^ In wild-type A549 cells, P-eGFP is observed to visually separate from M, indicative of successful fusion and content release **(Fig 3C, S3A, C)**. By contrast, in CD164^KO^ cells, P-eGFP and M remain colocalized in puncta, consistent with the presence of intact viral particles that fail to undergo fusion **(Fig 3C, S3A, C)**. Quantification of the extent of co-localization of P and M showed a profound fusion defect in CD164^KO^ cells for VSV-PICV **(Fig 3D)**. Infection by wild-type VSV was unaffected, indicating that the requirement is specific to arenavirus GPCs **(Fig 3D, Fig S3B)**.

### Clade A NWA GP1 binds directly to CD164 at acidic pH

To determine whether CD164 acts as a direct binding partner for GP1, we expressed and purified soluble GP1 proteins (sGP1) from PICV, PARV, FLEXV, and MACV, and soluble CD164 (sCD164) and the CD164 CRD (sCRD) (**Fig S4A**). Using bio-layer interferometry (BLI) we detected binding between PICV sGP1 and sCD164 at acidic pH (≤6.0), but not at neutral pH (**Fig 4A**). In addition, the sGP1 of PICV, PARV, and FLEXV bound sCD164 in a pH-dependent manner, whereas sGP1 of MACV failed to bind sCD164 regardless of pH (**Fig 4B, Fig S4B**). As expected, MACV sGP1 bound its receptor, human TfR1 (hTfR1), with a K_D_ of 2.52 μM ± 0.006 µM at pH=7.4 (**Fig S4D**).^43–46^ A multi-step kinetic analysis, however, demonstrated that MACV sGP1 does not bind hTfR1 at acidic pH (**Fig S4C**), whereas hTfR1 retains binding to a monoclonal anti-human CD71 antibody. This result indicates that MACV GP1 dissociates from its receptor during acidification of endosomes.

**Figure 4:**
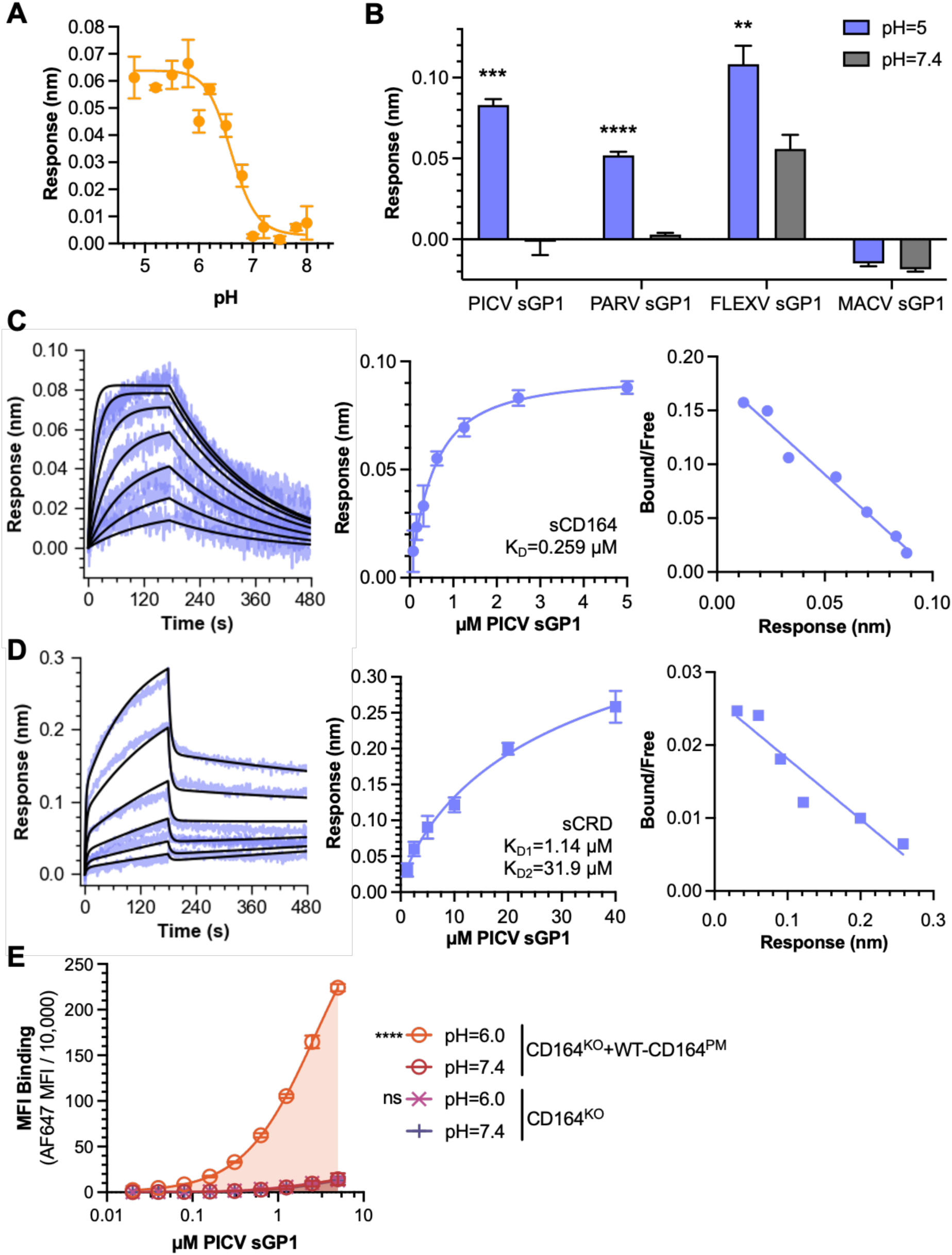
Effect of pH on the GP1-CD164 interaction. **(A)** Binding of 2.5 µM PICV sGP1 to 100 nM immobilized sCD164 as a function of pH; baseline corrected to binding of MACV sGP1 at same pH (n=3, independent experiments in triplicate). **(B)** Final binding response of 2.5 µM PICV, PARV, FLEXV, and MACV sGP1 to sCD164 at pH 5.0 (blue) or 7.4 (grey) (n=3, independent experiments in triplicate). One-Way ANOVA: \**p*<0.05, \*\**p*<0.005, \*\*\**p*<0.0005, \*\*\*\**p*<0.00005. **(C)** pH 5.0 binding curve of PICV sGP1 to sCD164 (100 nM). Left: 1:1 Langmuir fit; middle: nonlinear fit; right: Scatchard plot. Binding affinity (K_D_) calculated from 1:1 Langmuir fit, K_D_ = 0.259 µM ± 0.001 µM. (n=3, independent experiments in triplicate). **(D)** pH 5.0 binding of PICV sGP1 to soluble sCRD (200 nM). Left: 1:2 Langmuir fit; middle: nonlinear fit; right: Scatchard plot. Binding affinities (K_D_) calculated from 1:2 Langmuir fit K_D1_ = 1.144 μM ± 0.019 µM, and K_D2_ = 31.86 μM ± 0.555 µM. (n=3, independent experiments in triplicate) **(E)** Mean fluorescence intensity (MFI/10,000) of AF647-PICV sGP1 binding to CD164^KO^+WT-CD164^PM^ or CD164^KO^ cells at pH 6.0 or 7.4 (n=3, independent experiments in triplicate). One-Way ANOVA of AUC: *\*p<*0.05, *\*\*p<*0.005, *\*\*\*p<*0.0005, *\*\*\*\*p<*0.00005.

Kinetic analysis at pH 5.0 revealed that PICV sGP1 binds sCD164 with a K_D_ of 0.259 µM ± 0.001 µM (**Fig 4C**). Soluble PICV GP1 also bound directly to the sCRD in an acid pH-dependent manner but exhibited different kinetics compared to binding of sCD164 (**Fig 4D**). Binding of sGP1 to sCRD showed two kinetic phases consistent with a 1:2 Langmuir model characterized by the double-ascent and double-descent of the association and dissociation curves (**Fig 4D**). The slow-on/slow-off phase resembles binding of sGP1 to sCD164 with a K_D_ of 1.14 μM ± 0.019 µM, and the fast-on/fast-off phase may reflect reduced stabilization of the sCRD in the absence of MD2 with a K_D_ of 31.9 μM ± 0.555 µM (**Fig 4D**).

In cell-binding assays, Alexa Fluor 647 fluorescently labeled PICV sGP1 failed to bind CD164^KO^ cells but readily bound plasma-membrane targeted CD164-complemented cells at acidic pH (**Fig 4E**, **Fig S4E**). sGP1 bound to cells at acidic pH dissociated when cells were subsequently exposed to neutral pH (**Fig S4F)**. These observations support a model in which clade A NWA GP1 and CD164 interact directly at the pH found in acidified endolysosomal compartments.

### The C-terminus of the CD164 CRD mediates GP1 binding

AlphaFold3 modeling predicts that a β-sheet in PICV GP1 (residues 213-218) pairs with the C-terminal region of the CD164 CRD (residues 99-109) via main-chain hydrogen bonds (ipTM = 0.81, **Fig 5A**).^48–50^ To test this model, we performed deep mutational scanning (DMS) of the CRD displayed at the plasma membrane and sorted cells for binding of AF647-labeled PICV sGP1 at pH 6 (**Fig S5A**). This analysis identified that mutations in the C-terminal residues of the CRD were most detrimental to sGP1 binding (**Fig 5B, S5B**).

**Figure 5:**
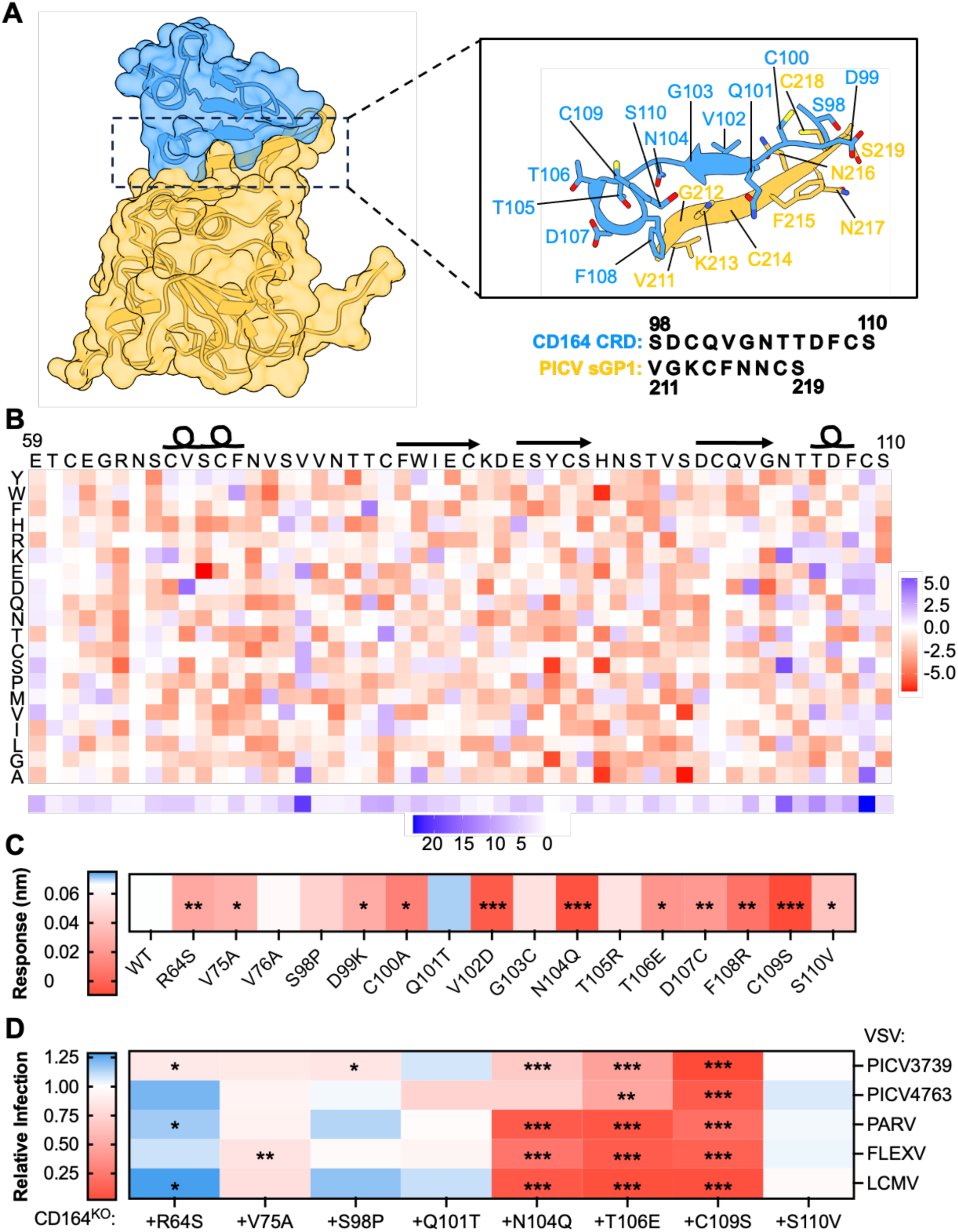
Requirements of the CD164 CRD for GPC-mediated binding and infection. **(A)** Alphafold3 predicted structure of PICV sGP1 (yellow) interacting with the CRD of CD164 (blue). Left: full predicted structure; right: zoom on predicted interaction region. Interface Predicted Template Modeling (ipTM) score=0.81. **(B)** DMS of CRD identifies key determinants for the acid-dependent GP1 interaction. DMS selection for loss of GP1 binding at pH 6.0 (GP1-sorted population S5A). Top: heat map of mutation differential scores (dms_tools2); blue: enriched mutations (>0), red: de-enriched (<0). Bottom: total enriched positive changes per amino acid site (score>0). (n=1, experimental replicate). **(C)** Averaged final association response of PICV sGP1 binding to sCD164 containing indicated mutations in CRD. Blue: response >WT sCD164 (0.066, white); red: response <WT. (n=3, independent experiments in triplicate). One-Way ANOVA: *\*p<*0.05, *\*\*p<*0.005, *\*\*\*p<*0.0005, *\*\*\*\*p<*0.00005. **(D)** CD164^KO^ cells expressing indicated CRD mutants were inoculated with chimeric VSV expressing indicated glycoproteins (MOI=1). % GFP-positive cells measured at 6 hours post-infection; normalized to WT HeLa (n=3, independent experiments in triplicate and technical replicates in duplicate). Blue: relative infection >WT (white, 1.0); red: <WT. One-Way ANOVA: *\*p<*0.05, *\*\*p<*0.005, *\*\*\*p<*0.0005, *\*\*\*\*p<*0.00005.

From this DMS analysis, we selected 16 CRD substitutions at either the distal (R64S, V75A, V76A) or proximal (S98P, D99K, C100A, Q101T, V102D, G103C, N104Q, T105R, T106E, D107C, F108R, C109S, S110V) interface that had the greatest effect on GP1 binding. We generated and purified sCD164 mutants (**Fig S6A**), tested their ability to bind a conformation-specific monoclonal antibody N6B6 as a proxy for CRD folding (**Fig S6B**), and examined their pH-dependent binding to PICV sGP1 (**Fig 5C**). Some mutations, including C100A, V102D, F108R and C109S disrupted recognition by an anti-CD164 antibody (N6B6) (**Fig S6B**), which may explain their failure to bind sGP1 (**Fig 5C**). Substitutions R64S, V75A, V76A, S98P, D99K, Q101T, N104Q, T105R, T106E, and S110V bind N6B6, and G103C and D107C retained partial binding by monoclonal antibody N6B6, indicative of preserved CRD folding. Of these mutations, the majority reduced binding to PICV sGP1 (**Fig 5C**), with N104Q showing the largest reduction, and substitutions R64S, V75A, S98P, D99K, G103C, T105R, T106E, D107C and S110V also reducing binding. Mutants V76A and Q101T were either unaffected in binding or had enhanced binding to sGP1 (**Fig 5C**). This analysis identifies residue N104 and the correct folding of the C-terminus of the CRD are required for recognition by sGP1.

To examine the consequence of specific mutations within CD164 for infection, we generated addbacks of R64S, V75A, S98P, Q101T, N104Q, T106E, C109S and S110V versions of CD164 in CD164^KO^ cells. Each of the mutants was expressed in cells, although the levels of Q101T were reduced (**Fig S6 C,D**). In contrast to our inability to detect C109S by monoclonal antibody N6B6 (**Fig S6B**), we detected its expression using polyclonal antisera (**Fig S6D, E**). Cells expressing mutations in the C-terminus of the CRD of CD164 were less susceptible to infection by VSV-PICV, VSV-PARV, VSV-FLEXV and VSV-LCMV, with C109S showing the largest inhibition (**Fig 5D**). The effect of any given substitution on infection was not uniform across clade A NWA VSV-chimeras (**Fig 5D**), suggesting virus-specific differences in their interaction with CD164. Consistent with this idea, the monoclonal anti-human CD164 antibody N6B6, which has been shown to inhibit LCMV, ^28^ and inhibits VSV-LCMV, does not inhibit infection by VSV-PICV or VSV-FLEXV and only partially impedes VSV-PARV (**Fig S6C**).^27^ Collectively, the binding and infection data support a model in which the C-terminus of the CRD forms main-chain mediated interactions that are highly sensitive to perturbation, linking local structural integrity to viral receptor engagement and function.

### A conserved β-sheet in GP1 mediates interaction with CD164

To further define the requirements for GP1 binding to CD164, we targeted the β-sheet of GP1 predicted to interact with CD164. We generated mutations in PICV sGP1 and assessed binding to sCD164 by BLI. For this analysis, we engineered mutations within the ß-sheet based on the predicted structure including: K213A, and double mutants K213A/C214A, F215A/N216A, N217A/C218A (**Fig S7A**). The double mutant F215A/N216A showed reduced binding to CD164, whereas K213A had no effect on binding, and substitutions to either cysteine (C214 or C218) destabilized sGP1 and abolished sCD164 binding (**Fig S7B, 6A**). When the same mutations were engineered into an infectious molecular clone of VSV-PICV3739, the cysteine mutations failed to yield infectious virus, consistent with the structural importance of this region. By contrast, we recovered virus containing substitutions K213A and the double mutant F215A/N216A. The double mutant F215A-N216A was able to infect CD164^KO^ cells, whereas the K213A mutant retained the requirement for CD164 for infection (**Fig 6B**).

**Figure 6:**
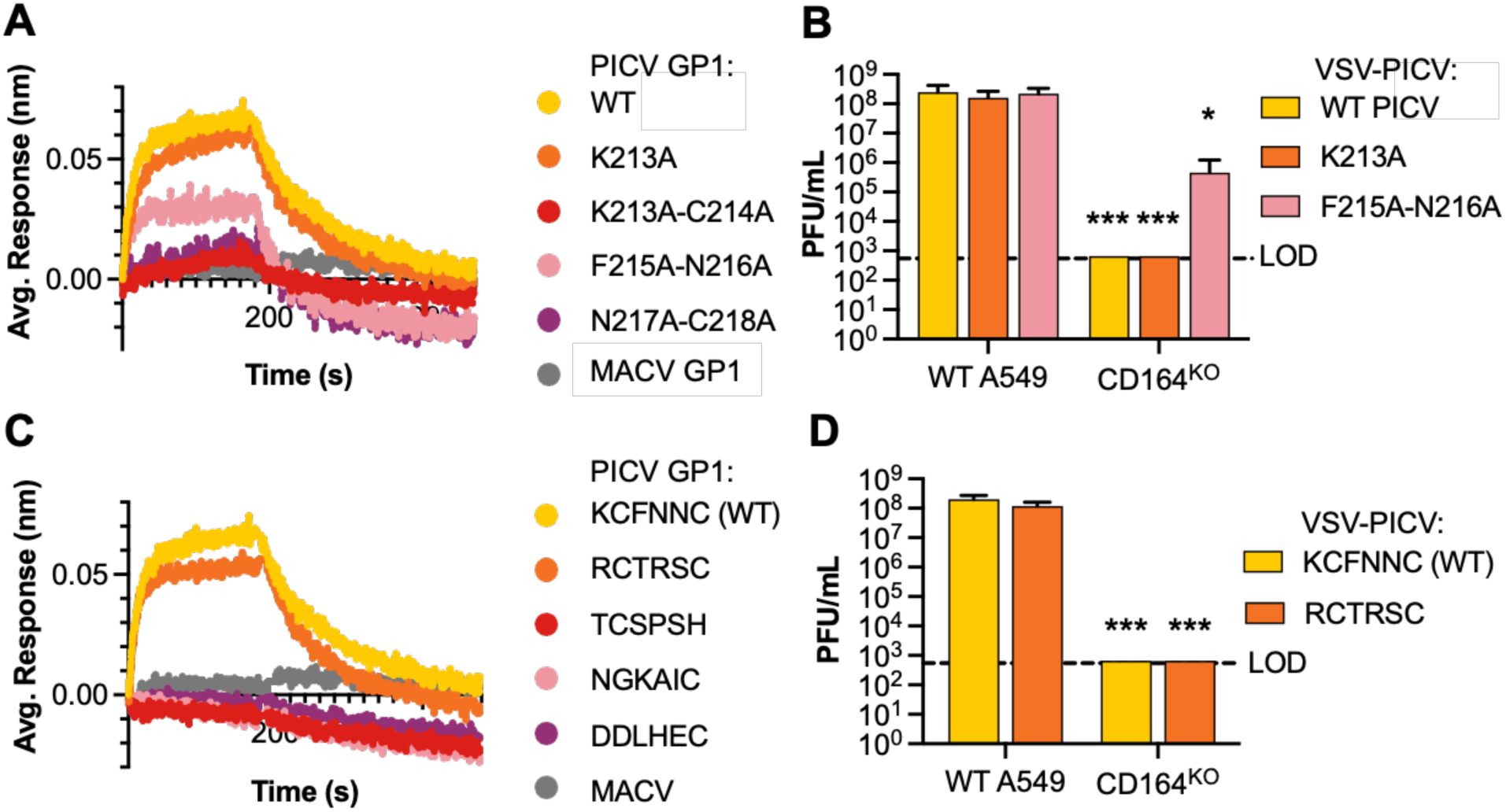
Analysis of the PICV GP1-CD164 interaction. **(A)** Impact of alanine mutations within the predicted ß-sheet of PICV sGP1 on binding to CD164. Binding of 2.5 µM WT or mutant PICV sGP1 to 100 nM of immobilized sCD164. Averaged response over association and dissociation measured by BLI. (n=3, independent experiments in triplicate). **(B)** Plaque assays in WT and CD164^KO^ A549 infected with VSV expressing WT PICV GPC (yellow), K213A mutant (orange), F215A-N216A mutant (pink). Dotted line: LOD. (n=3, independent experiments in triplicate). Two-tailed unpaired t-test, of CD164^KO^ compared to WT infected with indicated virus: *\*p<*0.05, *\*\*p<*0.005, *\*\*\*p<*0.0005. **(C)** Impact of stabilizing ß-sheet mutations (predicted by Protein MPNN) on CD164 binding, measured by BLI as in (A) (n=3, independent experiments in triplicate). **(D)** Impact of new consensus sequence generated by Protein MPNN analysis (RCTRSC, orange) on GPC-mediated infection by plaque assay in WT and CD164^KO^ A549 cells as in (B) (n=3, independent experiments in triplicate). Two-tailed unpaired t-test, of CD164^KO^ compared to WT infected with indicated virus: *\*p<*0.05, *\*\*p<*0.005, *\*\*\*p<*0.0005

Using ProteinMPNN we designed additional mutations to determine whether side-chain identity or β-sheet structure was critical for the interaction between GP1 and CD164. This yielded a consensus sequence (RCTRSC) in which the two cysteines and a predicted β-sheet were maintained while mutating all additional residues at the interface. A PICV sGP1 bearing this sequence expressed well, was stable, and bound sCD164 at acidic pH, whereas mutation of either cysteine was unstable and ablated binding (**Fig 6C, S7C, S7D**). A chimeric VSV-PICV with the RCTRSC mutation was infectious and retained CD164 dependence (**Fig 6D**). The predicted β-sheet is highly conserved across clade A NWA GP1 proteins with two conserved cysteines flanked by a lysine and a serine (**Fig S7E**). These findings indicate that the β-sheet structure – maintained by conserved cysteines – is required for the GP1 interaction with CD164 and subsequent infection, supporting a model in which main-chain hydrogen bonding rather than side-chain specificity mediates the GP1-CRD interaction.

## Discussion

In this study, we identify the endolysosomal sialomucin CD164 as a host-factor required for pH-dependent membrane fusion and cell entry of multiple clade A NWA. Using complementary genetic, biochemical, structural modeling and cell-based approaches, we define that CD164 functions as an endolysosomal receptor, extending the receptor-switching mechanism previously defined for OWA.^26–29^ Although CD164 is transiently present on the plasma membrane, we find that its interaction with clade A NWA GP1 is strictly dependent upon acidic pH. This pH dependence, coupled with the predominant endolysosomal localization of CD164, supports a model of clade A NWA infection that depends on engagement of yet-to-be-identified attachment factor(s) at the cell surface with a subsequent switch to bind CD164 in the acidic environment of endosomes for membrane fusion. This mirrors the two-receptor entry pathway established for LASV, LCMV, and LUJV, all of which rely on plasma membrane attachment followed by pH-induced engagement of LAMP1, CD164, or CD63, respectively.^26–29^ Our data therefore suggest that a unifying feature of arenavirus entry is the requirement for endolysosomal receptors and acidic pH to trigger GP2-dependent fusion. Fusion assays further support a sequential, pH-coordinated entry mechanism. CD164 engagement occurs at higher pH than GPC fusion (≤6.0 versus 5.0–5.4), suggesting a multi-step process in which GP1 binds CD164 in late endosomes, promoting GP1 destabilization and priming for GP2-mediated fusion.

The clade A NWA have converged on the same endolysosomal receptor as LCMV despite substantial sequence divergence. PICV GPC shares only 43% identity with LCMV GPC, yet both utilize CD164. However, mutations within the CD164 CRD, and inhibition of engagement by an antibody, have distinct effects on clade A NWA infection and LCMV infection, indicative of different engagement of this shared factor. We note that LASV and LCMV which share 61% identity and share a plasma membrane receptor (α-DG) diverge in their endolysosomal fusion receptors. These contrasting patterns highlight the flexibility in GP1-host interactions and suggest that endolysosomal receptor usage is a conserved feature rather than a lineage-specific adaptation. Our structure-function analyses further define the molecular basis of the GP1-CD164 interaction. Deep mutational scanning, targeted mutagenesis, and AlphaFold3 modeling converge to show that the C-terminus of the CD164 CRD interacts with a conserved β-sheet within the clade A NWA GP1. The interface is dominated by main-chain hydrogen bonding rather than side-chain specificity, which explains both the broad tolerance for some mutations and the strict requirement for correct CRD folding. Likewise, the essential role of conserved cysteines within the GP1 β-sheet suggests that structural integrity is required for productive interaction. This interaction mode parallels other viral spike-receptor interfaces, such as HIV-1 gp120 and CD4, in which β-sheet pairing and main-chain hydrogen bonding dominate binding, underscoring how backbone-mediated contacts can enforce structural precision during entry.^51,52^

Comparison of biochemical binding and infection phenotypes provides insight into the functional architecture of the CRD. Mutations such as N104Q and T106E impaired both sGP1 binding and viral entry, supporting a direct role in the predicted main-chain mediated GP1 interface. In contrast, some mutations displayed disproportionate effects. For example, R64S reduced sGP1 binding yet produced more modest defects, or even enhancement, in infection, suggesting that avidity effects, membrane context, or additional regions of CD164 are of greater influence during infection. Conversely, C109S caused a pronounced infection defect across multiple arenaviruses consistent with heightened sensitivity of viral entry to subtle structural destabilization or requirements beyond attachment. Together, these findings indicate that while sGP1 binding predicts receptor engagement, productive infection imposes additional structural constraints at the CRD C-terminus, defining a conserved yet flexible finely tuned mechanism by which clade A NWA engage CD164. The finding that the Machupo virus GP1-transferrin receptor interaction is acid labile suggests that this receptor switching paradigm may extend to the clade B NWA. However, this observation contrasts with a recent report describing MACV GP1 binding to a rodent TfR1 ortholog at acidic pH in a single-step kinetic analysis.^53^ We note however, that study relied upon an engineered apical domain of *Neotoma albigula* TfR1 designed for broad antiviral immunotherapeutic applications to remain bound to GPC and act as an inhibitor for entry, rather than for mechanistic interrogation of receptor engagement.^53,54^

An important limitation of our work is the absence of identified plasma-membrane attachment factors for clade A NWA. This precludes a full reconstruction of the sequential entry pathway. The identification of factors that are bound at the plasma membrane is necessary as receptor switching relies on the coordinated handoff between binding of surface factors and endolysosomal factors for the OWA. Structural studies of clade A NWA GPC are also needed to resolve the predicted GP1 β-sheet. An additional limitation of this study includes the inability to assess the impact of CD164 in pathogenesis *in vivo*. Our observation that the antibody, N6B6, does not block clade A NWA infection in *in vitro* suggests we would be unable to effectively prevent CD164-mediated pathogenesis in guinea pig^55^ or hamster models.^56^ In addition, PICV does not produce tractable disease or pathogenic phenotypes in mice, which is the only system with a conditional CD164 knockout currently available. Consequently the impact of CD164 expression on pathogenesis was not evaluated in this study.^57^

Together, our findings establish CD164 as an endolysosomal receptor for a NWA and reveal mechanistic parallels in receptor engagement and fusion across *Arenaviridae*. These insights expand our understanding of arenavirus entry biology and suggest that endolysosomal receptor switching is a conserved and unifying feature of mammalian arenaviruses.

## Acknowledgements

We thank members of the Whelan laboratory for helpful discussions and critical feedback throughout the course of this study. We thank Wandy Beatty and the Molecular Microbiology Imaging Facility at Washington University in St. Louis for assistance with transmission electron microscopy, and the Department of Pathology and Immunology Flow Cytometry and Fluorescence Cell Sorting Core for assistance with cell sorting. We thank the World Reference Center for Emerging Viruses and Arboviruses (WRCEVA) for providing the Pichindé virus strain used in this study. We thank the Washington University Center for Cellular Imaging (WUCCI) for imaging using the Zeiss LSM 980 Airyscan Confocal Microscopy. We thank the Genome Access Technology Center (GTAC) at the McDonnell Genome Institute at Washington University School of Medicine for Illumina Miseq sequencing; and the Genome Engineering and iPSV core (GEiC) at Washington University School of Medicine for development of the plasma membrane targeted loss-of-function CRISPR plasmid library.

This work was supported in part by National Institutes of Health grant **U19AI81984** (to S.P.J.W). C.E.T. was supported in part by training grants **T32GM007067** and **T32GM139774** (to Heather True-Knob).

## Author Contributions

C.E.T. and S.P.J.W. designed the study; C.E.T. conducted all experiments resulting in published data. T.K. played key roles in BLI development and analysis. R.M.M. analyzed CRD DMS data. M.C.P. performed microscopy and imaging analysis. P.W.R. conducted MAGeCK analysis of sgRNA enrichments from CRISPR screens. Z.L., J.D., and H.M. developed CRISPR libraries. M.A.R. provided experimental support generating PICV mutants. C.E.T. and S.P.J.W. wrote the intial draft with all authors providing comments. Supervision and funding acquisition was caried out by S.P.J.W., D.H.F., and M.S.D.

---

**Supplemental Figure 1: Related to Figure 1:**
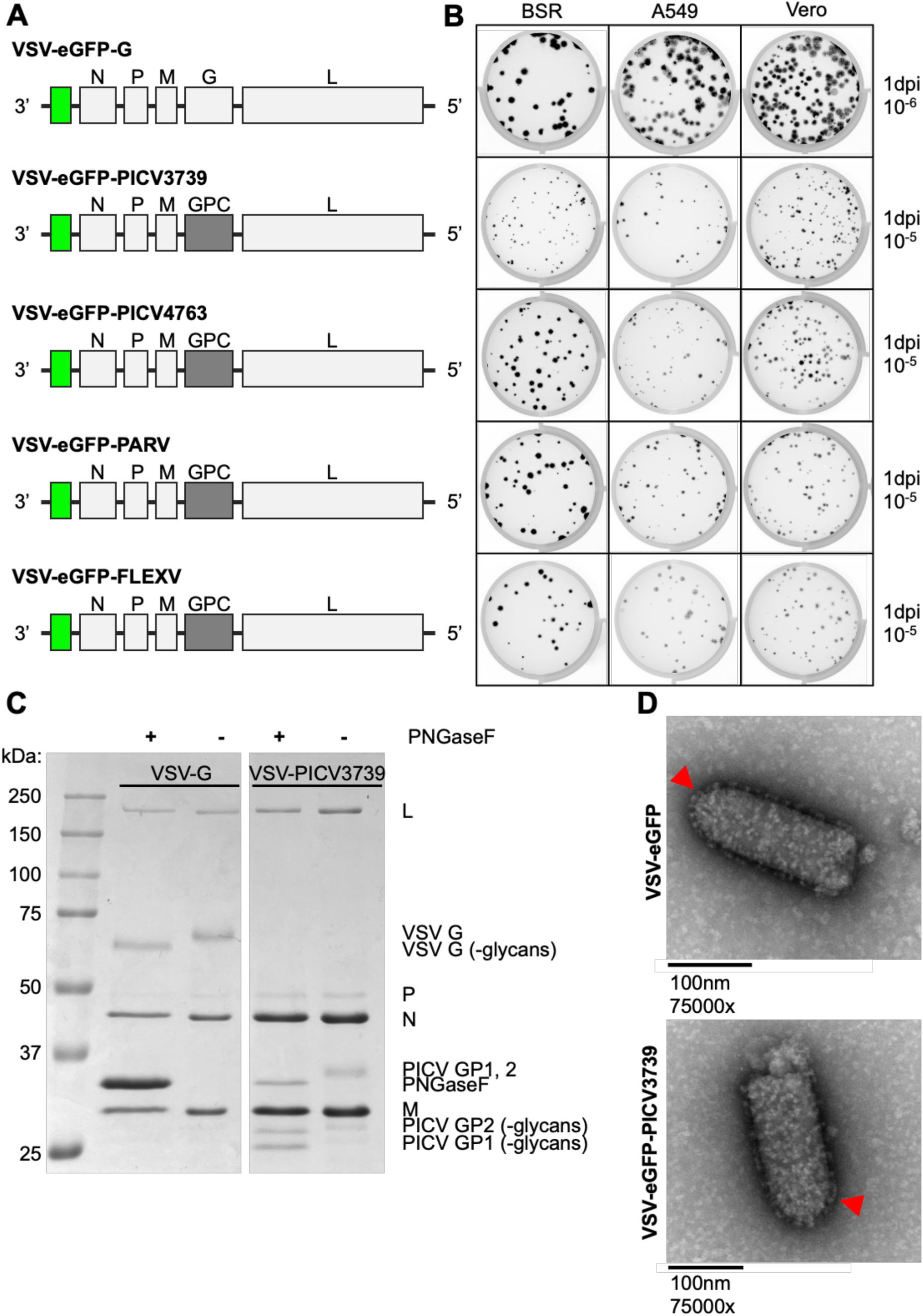
Chimeric VSV expressing clade A New World arenavirus GPC. **(A)** Schematic of chimeric VSV genomes expressing NW arenavirus glycoproteins (VSV-eGFP-NW-GPC; PICV CoAN4763/3739, PARV, FLEXV) in place of native VSV G and expressing eGFP as infection marker. Gene order 3’→5’: eGFP (green), N, P, M, G (light grey) or NW-GPC (dark grey), L. **(B)** Focus-forming unit assays at 1 dpi of indicated cells infected with chimeric VSVs. Representative wells: 10^-6^ (VSV-eGFP-G) or 10^-5^ (VSV-eGFP-NW-GPC) dilutions. **(C)** Coomassie-stained SDS-PAGE of purified VSV-eGFP-G or VSV-eGFP-PICV CoAN3739 (VSV-PICV3739), ± PNGaseF treatment. **(D)** Representative image from negative-stain electron microscopy of gradient purified VSV-eGFP-G or VSV-PICV3739. Scale bar=100 nm. Red arrows indicate virion surface glycoproteins.

**Supplemental Figure 2: Related to Figure 1:**
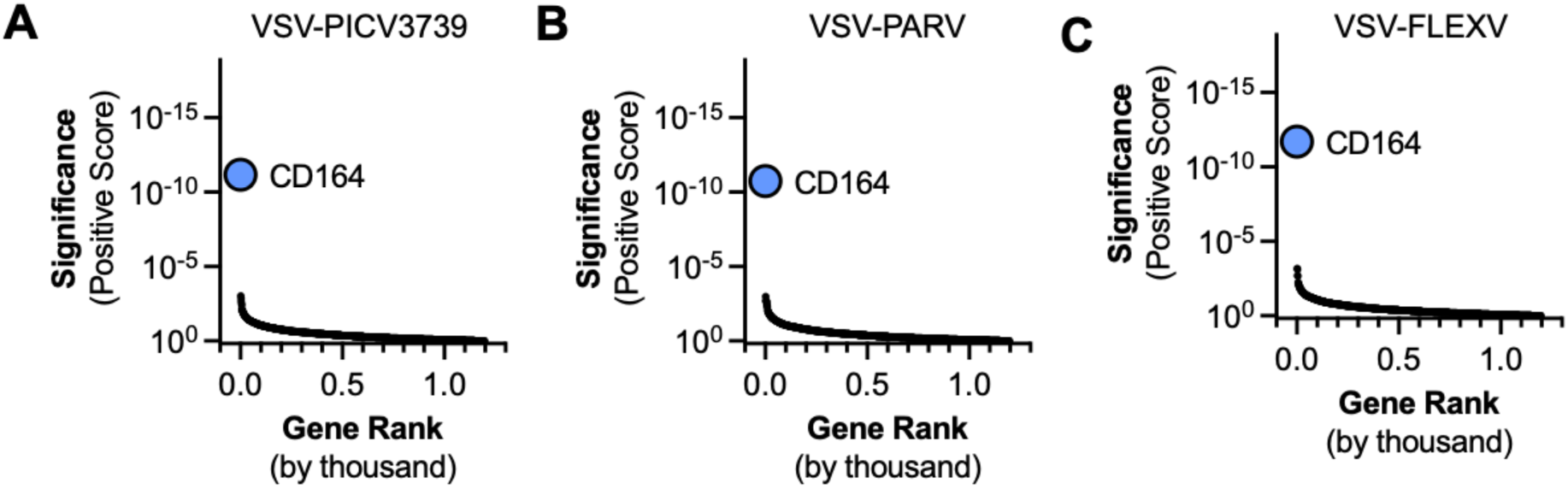
Genetic screen of surfaceome proteins required for clade A NWA infection. Enrichment of guide RNAs from a CRISPR-Cas9 HAP1 plasma membrane protein (surfaceome) screen after one round of infection with chimeric **(A)** VSV-eGFP-PICV (CoAN3739) **(B)** VSV-PARV **(C)** VSV-FLEXV at an MOI of 3. Genes arranged by rank on the x-axis; significance (positive score) on the y-axis. (n=1, experimental replicate).

**Supplemental Figure 3: Related to Figure 3:**
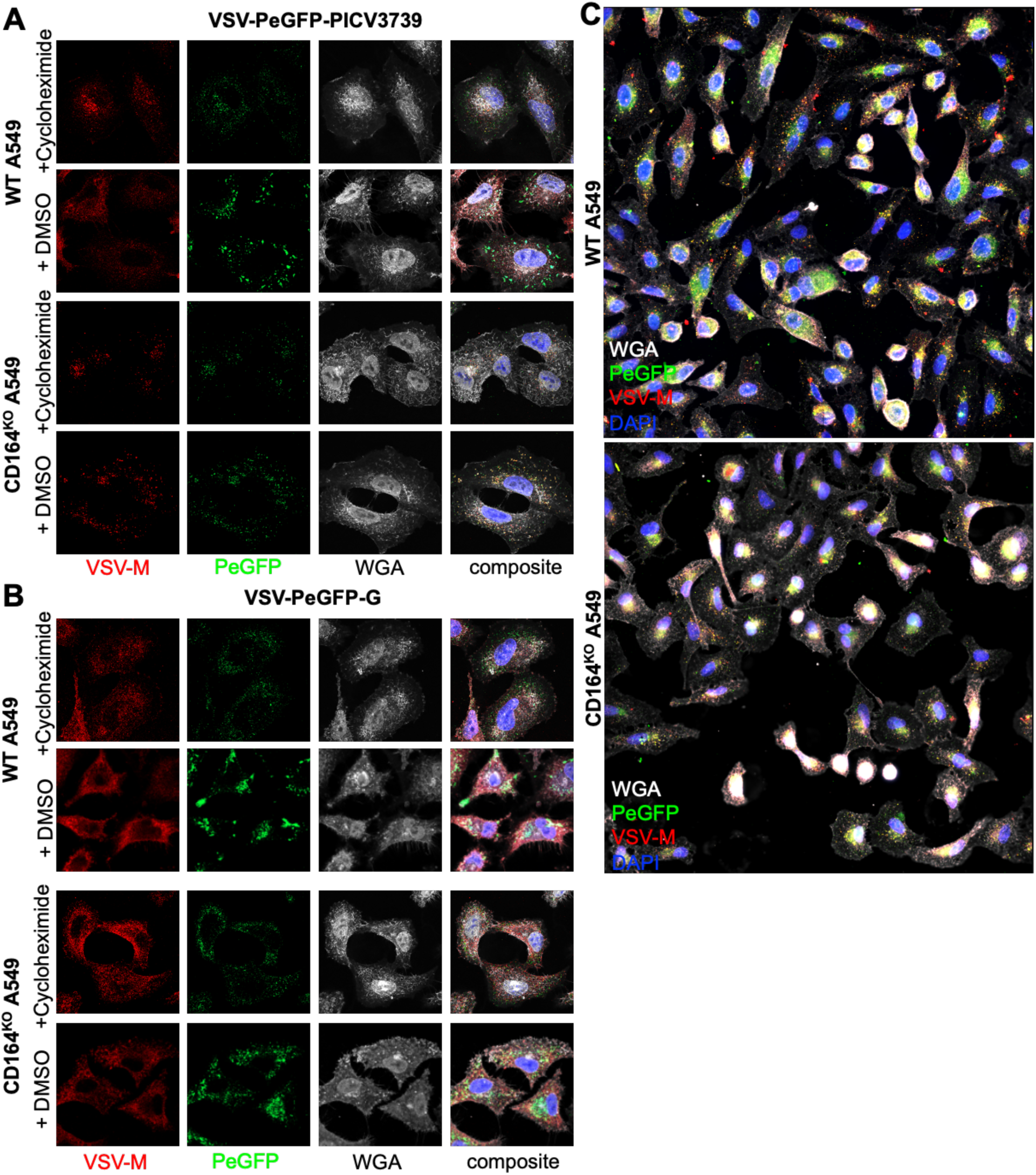
Effect of CD164 on GPC-mediated fusion. (A-B) Additional representative images from Figure 3C. WT or CD164^KO^ A549 cells pretreated with cycloheximide or DMSO, inoculated with VSV-P/eGFP-PICV3739 (A, 3 hr) or VSV-P/eGFP-G (B, 3 h). Cells were fixed, stained with DAPI (blue), WGA (Alexa 647, white), anti-VSV-M (Alexa 594, red), imaged by Airyscan confocal microscopy. Scale bar, 10 µm. **(C)** Widefield microscopy of WT or CD164^KO^ A549 pretreated with cycloheximide, inoculated with VSV-P/eGFP-PICV3739, fixed, and stained at 2 hours post-infection.

**Supplemental Figure 4: Related to Figure 4.**
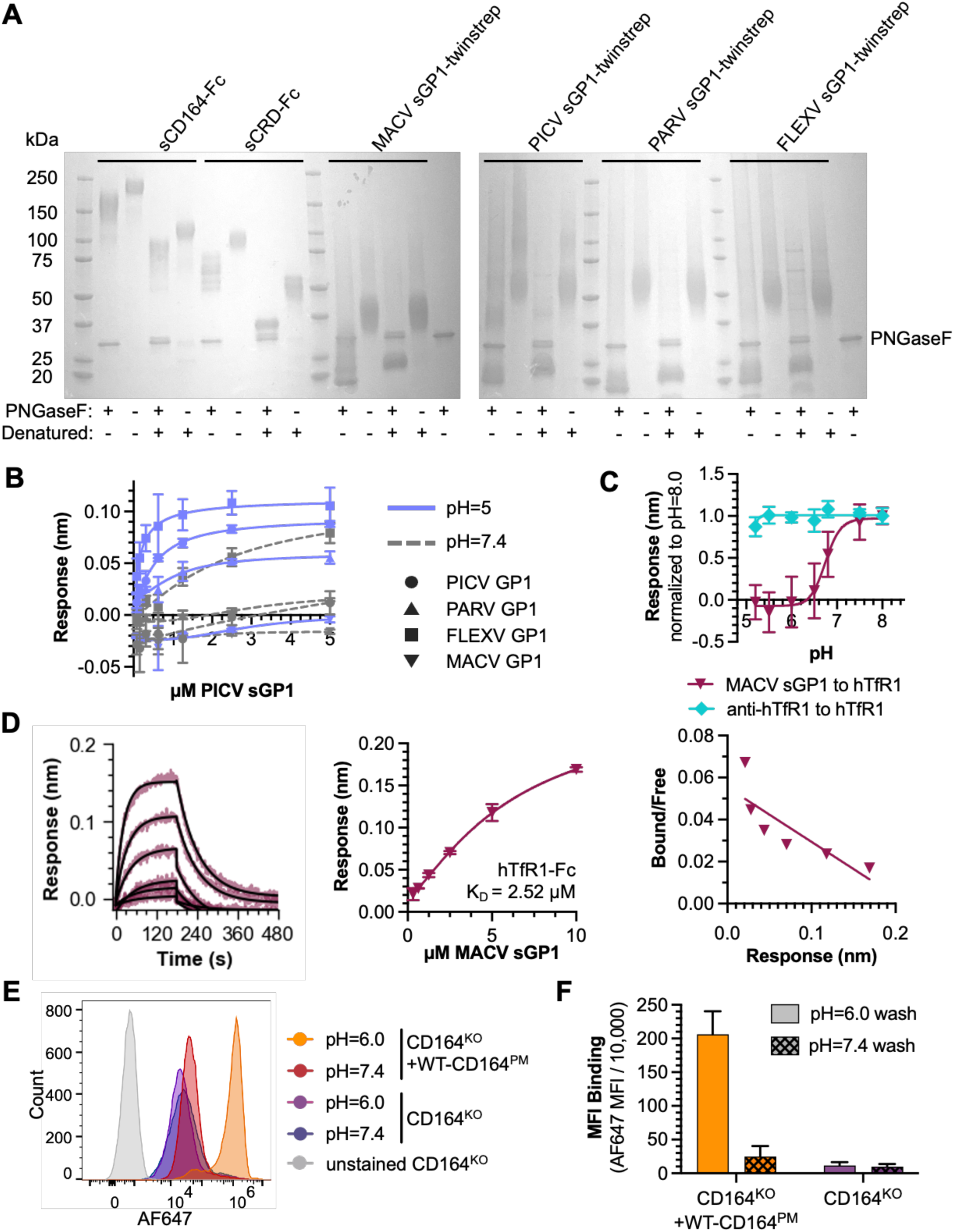
Effect of pH on the GP1-receptor interaction. **(A)** Coomassie-stained SDS-PAGE of soluble proteins used in Figure 4A-E. 2.5 µg of PNGaseF treated, nondenatured, or denatured purified protein. **(B)** Nonlinear fit of binding curves at pH 5.0 (blue) and 7.4 (grey) for PICV sGP1, PARV sGP1, FLEXV sGP1, and MACV sGP1 to 100 nM of immobilized sCD164. Curves represent the averaged final response at the end of association (n=3, independent experiments in triplicate). **(C)** Binding of 5 µM MACV sGP1 (maroon triangle) or 1 µg mouse anti-human TfR1 (CD71) antibody (cyan diamond) to 100 nM of immobilized hTfR1-Fc as a function of pH. Normalized to max binding at pH 8.0 (n=2, independent experiments in duplicate). **(D)** pH 7.4 binding curve of MACV sGP1 to hTfR1 (100 nM). Left: 1:1 Langmuir fit; middle: nonlinear fit; right: Scatchard plot. Binding affinity (K_D_) calculated from 1:1 Langmuir fit, K_D_ = 2.52 µM ± 0.006 µM. (n=2, independent experiments in duplicate). **(E)** Representative histogram of mean fluorescence intensity (MFI) from Figure 4E. Cells bound with 2.5 µM AF647-labeled PICV sGP1 at pH=6.0 (orange: CD164^KO^+WT-CD164^PM^; purple: CD164^KO^) or pH 7.4 (red: CD164^KO^+CD164^PM^; blue: CD164^KO^) (n=3, independent experiments in triplicate) **(F)** MFI (/10,000) of CD164^KO^+WT-CD164^PM^ or CD164^KO^ cells from Figure 4E and S4E, labeled with 5 µM AF647-PICV sGP1 at pH 6.0 and then washed at pH 7.4 (hashed) or 6.0 (solid). (n=3, independent experiments in triplicate).

**Supplemental Figure 5: Related to Figure 5:**
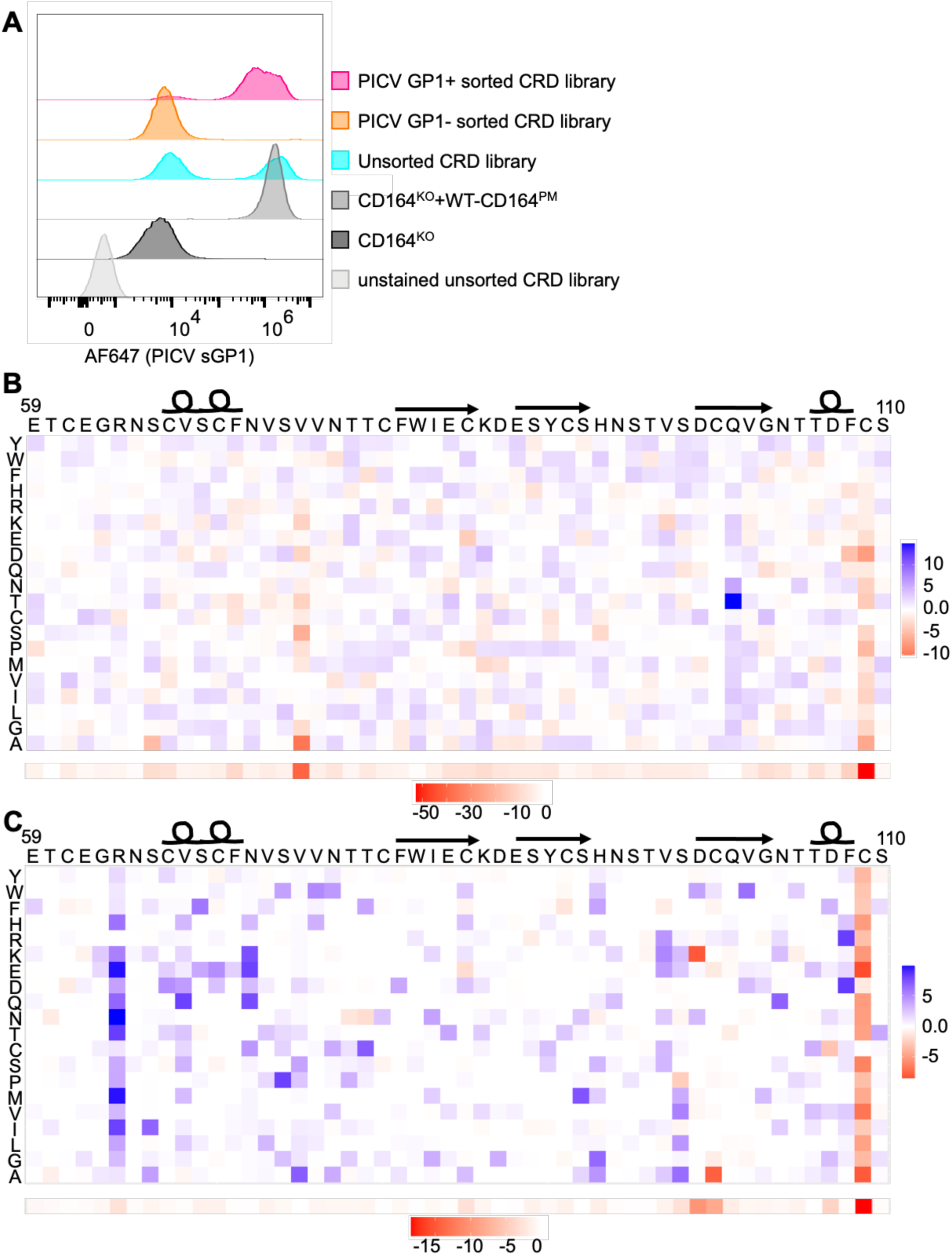
Deep mutational scanning (DMS) of the CRD of CD164. **(A)** Representative histograms of AF647-PICV sGP1 binding to CRD library (cyan) and resulting GP1-(orange) and GP1+ (pink) populations sorted from CRD library. AF647 fluorescence (x-axis) and count (y-axis). **(B-C)** DMS of CRD identifies key determinants for acid-dependent GP1 interaction and N6B6 monoclonal antibody binding. (B) DMS selection for GP1 binding at pH 6.0 (GP1+ sorted population). Top: heat map of mutation differential scores (dms_tools2); blue: enriched mutations (>0), red: de-enriched (<0). Bottom: total enriched changes per amino acid site (score>0). (C) DMS selection for N6B6 binding at pH 7.4, same representation as in (B). (n=1, experimental replicate).

**Supplemental Figure 6: Related to Figure 5:**
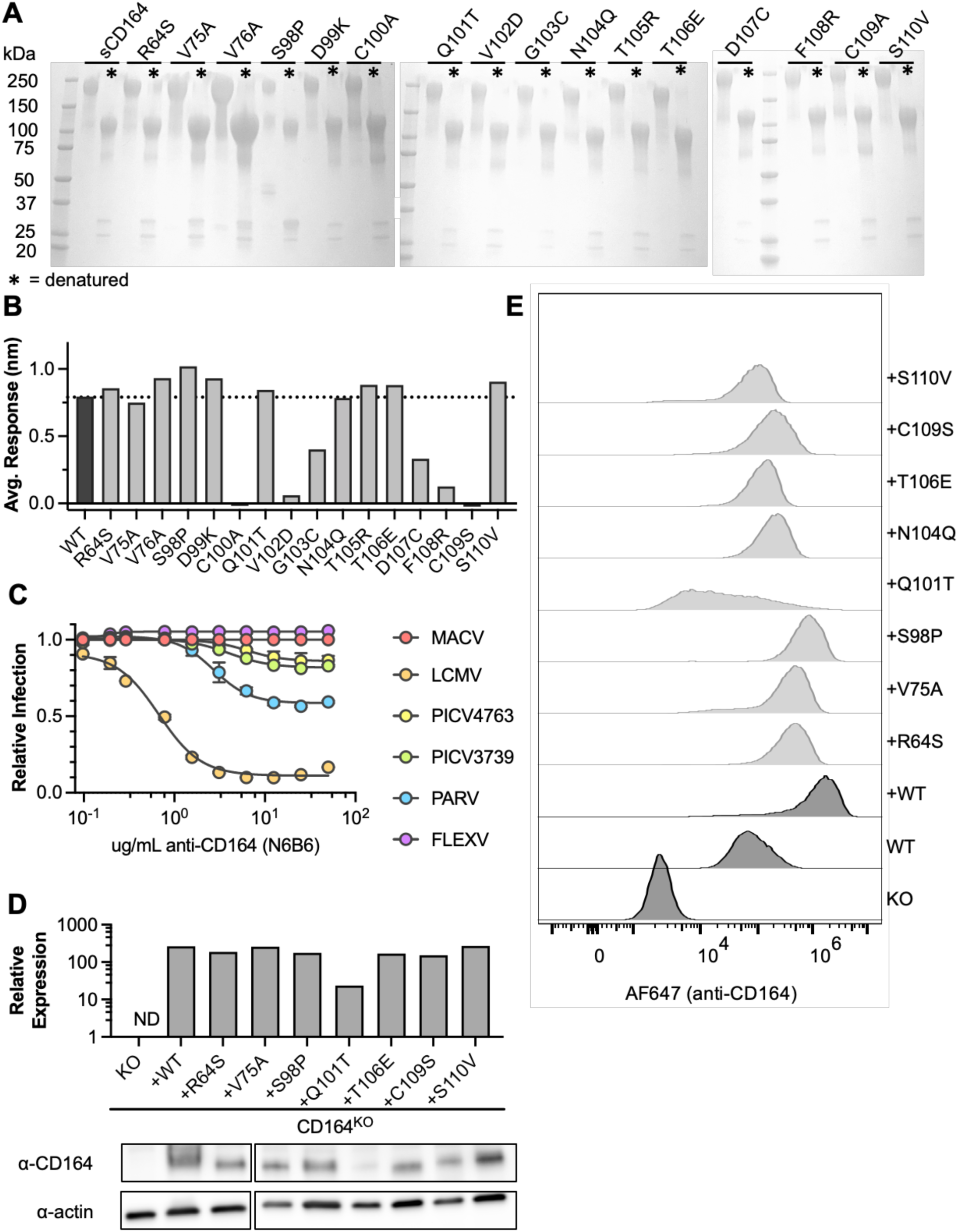
Deep mutational scanning of the CD164 CRD. **(A)** Reducing/denaturing or non-denaturing Coomassie of sCD164 mutant proteins (Figure 6B). Asterisk (*) denotes denaturing. **(B)** Final binding response of 1 µg N6B6 anti-human CD164 antibody to 100nM immobilized sCD164 mutants. **(C)** Inhibition of chimeric VSV expressing indicated glycoproteins by anti-CD164 monoclonal antibody, N6B6. Cells were pretreated with N6B6 for 1 hour on ice, virus was then added at MOI=1. At 6 hours post-infection, cells were fixed and % GFP-positive cells was determined by flow cytometry. % GPF+ was normalized to HeLa cells infected with no inhibition (n=2). **(D)** Relative expression of CD164 in CD164 mutant addback cells. Top: area under the curve (CD164/actin) compared to WT; bottom: representative Western blot. **(E)** Representative histograms of total CD164 expression in CD164 mutant addback cells.

**Supplemental Figure 7: Related to Figure 6:**
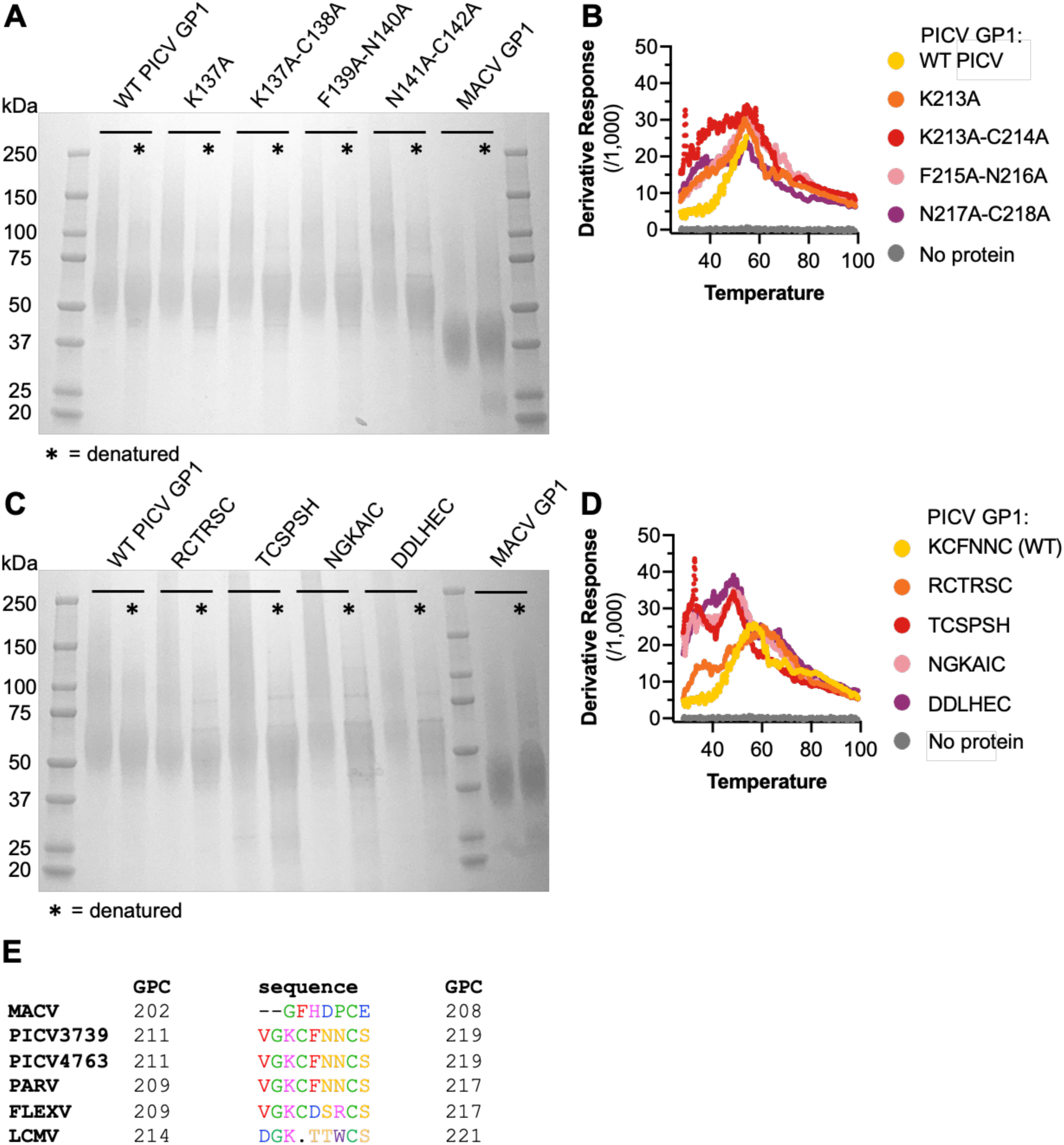
Analysis of the PICV GP1-CD164 interaction. **(A)** Reducing and denaturing or non-denaturing Coomassie of soluble PICV sGP1 containing alanine mutations (used in Figure 6A). Asterisk (*) denotes denaturing conditions. **(B)** Thermal stability shift assays of PICV sGP1 alanine mutants as a proxy for folding (in technical quadruplicate). **(C)** Reducing and denaturing and non-denaturing Coomassie of PICV sGP1 containing ß-sheet mutants (used in Figure 6C). Asterisk (*) denotes denaturing. **(D)** Thermal stability shift assays of PICV sGP1 ß-sheet mutants as a proxy for folding (in technical quadruplicate). **(E)** Sequence alignment of GPCs (MACV, PICV3739, PICV4763, PARV, FLEXV, and LCMV) in the predicted ß-sheet region of interaction. Alignment performed using Clustal Omega via EMBL-EBI; numbering refers to full-length GPC. Red indicating residues with hydrophobic side chains, green indicating special case residues, pink indicating residues with positively charged side chains, orange indicating residues with polar uncharged side chains, and blue indicating residues with negatively charged side chains.

## Methods

### Cells

Cell lines used in this study: A549 (a gift from Nir Hacohen, Broad Institute of Massachusetts Institute of Technology and Harvard, Cambridge, MA), HeLa (a gift from Jan Carette, Department of Microbiology and Immunology, Stanford University School of Medicine, Stanford, CA), HEK239T (ATCC No. CRL-32169), Vero CCL81 (ATCC CCL-81), Expi293™ (ThermoFisher Scientific, A14635), and BSRT7.^58^ All cell lines used in this study, including knockouts and addbacks, other than Expi293™ cells, were cultured in Dulbecco’s modified Eagle’s medium (DMEM, Corning, 10-013-CMR) supplemented with 10% fetal bovine serum (FBS) (Corning, 35-015-CV) and maintained at 37°C and 5% CO_2_. Expi293™ cells were cultured in Expi293 Expression Medium and maintained at 37°C, 8% CO_2_, and shaken at 125 rpm. Knockout cells (CD164^KO^) and many complemented addback cells (CD164^KO^: +CD164, +ΔMD1, +ΔCRD, +ΔMD2, +CRD-LAMP1, +WT-CD164^PM^, +N104Q-CD164) used in this study were generated previously.^27^

### Generation of lentiviral addback cell lines

A lentivirus delivery system was used to reconstitute protein expression in HeLa and A549 CD164^KO^ cells using a lentiviral cDNA expression plasmid, pCW62-Puro (Harvard plasmid repository No. EvNO00438621).^27^ Recombinant lentiviruses generated in this study (CD164 mutants, **Figure 5C, S6C-D**) were produced by PEIMax (Kyfora Bio, 24765) mediated transfection of HEK293T cells with pCW62 expressing the gene of interest (CD164 mutant), pCAGGS-VSV-G, and psPAX2. Cell supernatants were collected 48 hours post transfection and passed through a 0.45 μM filter. Addback cell lines generated in this study were produced by low MOI transduction via spinoculation in the presence of 10 μg/mL polybrene (TR-

1003-50UL) for 30 min at room temperature. Cells were maintained in DMEM + 10% FBS supplemented with 2 μg/mL of puromycin (Gibco, A1113803) starting 4 hours post spinoculation.

### Generation of Chimeric VSVs

Chimeric VSVs expressing eGFP in the first position and the GPCs of MACV (Carvallo strain^29^) and LCMV (Armstrong strain^16,29^) were generated previously. VSV-PICV3739 (GenBank: QLA46850.1), VSV-PICV4763 (Munchique, GenBank: ABU39904.1), VSV-PARV (GenBank: YP_001936017), and VSV-FLEXV (Pinheiro, GenBank: AAN09937.1) were generated in this study. Cloning and rescue of chimeric VSVs were performed as previously described.^29,59^ In brief, gene blocks containing the GPC sequences, purchased from Integrated DNA Technologies, were cloned into MluI and NotI digested VSV backbone (pVSV-eGFP-GPC). BSRT7 cells were infected with vaccinia virus (vTF7-3) at an multiplicity of infection (MOI) of 3 for 1 hr then transfected with Lipofectamine 2000 (Invitrogen, 11668027) and the pVSV-eGFP-NW-GPC plasmid along with pCAGGS helper plasmids encoding VSV-L, VSV-G, VSV-N, and VSV-P and incubated for 4-6 hr at 37°C. Media was then changed to DMEM supplemented with 2% (v/v) FBS and 20 mM HEPES (Sigma-Aldrich, H3375), and cells were incubated for an additional 3 days at 34°C and 5% CO_2_. Supernatants were harvested and clarified by centrifugation and filtration through a 0.22 μM filter. Processed rescue supernatants were titrated on BSRT7 cells in the presence of 40 μg/mL of cytosine arabinoside (AraC, Sigma-Aldrich, C1768) via plaque assay to determine rescue efficiency. Individual virus clones were isolated through agarose plugs and grown on BSRT7 cells in the presence of AraC to generate p1 stocks. RNA from infected cells were extracted using TRIzol Reagent (Invitrogen, 15596018). The glycoprotein (GPC) of interest was amplified from RNA by RT-PCR (Qiagen One-Step, QIAGEN, 210212) utilizing VSV specific primers upstream and downstream of GPC. cDNA was gel extracted and sequenced by Sanger sequencing. Working stocks were grown on BSRT7 cells at a MOI of 0.01 for 48-72 hr in DMEM + 2% (v/v) FBS + 20 mM HEPES. All chimeric VSVs were titrated by plaque assay and flow cytometry on A549, HeLa, and BSRT7.

### Wild-type arenavirus infections

Virus stocks of: LCMV (Armstrong LCMV-2A-GFP, a gift from Juan Carlos de la Torre, Scripps), PARV (12056, ATCC VR-477), and PICV4763 (a gift from the World Reference Center for Emerging Viruses and Arboviruses (WRCEVA), University of Texas Medical Branch (UTMB), Galveston, TX), were grown on BSRT7 cells at a an MOI of 0.01 for 5-7 days in DMEM supplemented with 2% (v/v) FBS and 20 mM HEPES. Stocks were titrated by plaque assay on BSRT7 and A549. Plaque assays were conducted by infection of A549 cells in decreasing 1:10 dilutions for 1 hour at 37°C. Viruses were then removed from cells and overlay media (MEM + 2% (v/v) FBS + 20 mM HEPES + 0.75% (v/v) NaHCO_3_ + 2 mM glutamine) supplemented with 0.25% (w/v) agarose (Sigma-Aldrich, A9539) was added and infection was allowed to proceed for 5 days at 37°C. Cells were then fixed in 2-5% (v/v) paraformaldehyde overnight at room temperature. Agarose was then removed, and cells were stained with 1% (w/v) crystal violet (ThermoFisher Scientific, B21932.36) and 20% (v/v) methanol (Sigma-Aldrich, 34860) for 1 hour at room temperature. Stain was removed and cells were washed with water, dried, and plaques were counted.

### Purification of Recombinant VSV

BSRT7 cells were infected at near confluency at an MOI of 3 in 5 mL of DMEM + 2% (v/v) FBS + 20 mM HEPES. Viruses were absorbed to cells for 1 hr at 37°C with gentle rocking every 10 min. Inoculum was removed and replaced with 15 mL of OptiMEM and infections were incubated at 34°C for 72 hr. Cell supernatants were then clarified by centrifugation at 3000 rcf for 10 min. Clarified supernatants were then transferred to polycarbonate 30 mL Oak Ridge Tubes (ThermoFisher Scientific, 3118-0030), and viruses were pelleted by centrifugation using a Beckman Coultier Optima XPN-100 Ultracentrifuge and Type 70Ti rotor at 41,692 rcf for 90 min at 4°C. Supernatants were then aspirated, and the virus pellet was resuspended overnight at 4°C on ice in 0.5 mL NTE (0.1 M NaCl, 1 mM EDTA, 0.01 M Tris, pH 7.4). Virus was then further purified using a 15-45% (w/v) sucrose gradient, utilizing Beckman Ultra-clear tubes (Beckman Coultier, 34405-9) in a SW-41 rotor, centrifuging for 3 hr at 150,920 rcf. The visible band of virus was then collected from the tube by side puncture in minimal volume and dialyzed (Slide-A-Lyzer Dialysis Cassette, ThermoFisher Scientific, 66005) into PBS (Quality Biological, 119-069-151) overnight. Protein was quantified using Bradford Reagent (Sigma Aldrish, B6916). 5 μg of gradient purified virus was treated with PNGaseF using manufacturer’s protocols (New England BioLabs, P0704) under denaturing conditions. PNGaseF treated and untreated samples were run on a 12% (w/v) acrylamide low bis SDS-PAGE gel under denaturing and reducing conditions and imaged by Transmission Electron Microscopy (TEM) to determine glycoprotein incorporation and quality.

### Transmission electron microscopy

Gradient purified VSV-eGFP-G and VSV-eGFP-PICV (3739) were imaged by Transmission Electron Microscopy (TEM) in collaboration with Wandy Beatty in the Molecular Microbiology Imaging Facility (Washington University School of Medicine, St. Louis). Prior to staining, glutaraldehyde was added to a final concentration of 1% (w/v) to virus stocks. Samples were then negative stained with 0.75% (w/v) uranyl formate (pH 7.5). Ten images were collected on a JEOL 1200 EX II Transmission Electron Microscope at 75,000X, displaying one representative image per sample.

### Genome-Wide Loss-of-Function CRISPR-Cas9 Library Generation

An African Green Monkey (*Chlorocebus sabaeus*) single guide RNA (sgRNA) knockout library was generated in Vero cells using a two-plasmid system. Vero cells were transduced with a lentiviral vector expressing SpCas9 and selected with 40 μg/mL blasticidin for one week. Blasticidin-resistant Vero-Cas9 cells were subsequently single-cell cloned, and individual clones were assessed for functional Cas9 activity using a GFP reporter assay.^60^ The African green monkey CRISPR lentiviral library contained 76,411 sgRNAs targeting 19,114 genes, along with 1,000 non-targeting control sgRNAs. To generate the pooled knockout population, 1.2x10^8^ Vero-Cas9 cells were transduced at a MOI of 0.3 with the African green monkey library and selected with puromycin (10 μg/mL) for 10 days. Surviving cells were expanded and maintained in DMEM + 10% (v/v) FBS and then stored in liquid nitrogen. Library representation and sgRNA coverage was assessed by sequencing genomic DNA.

### Genome-Wide Loss-of-Function CRISPR-Cas9 Screens

For screening, library cells were infected with chimeric VSV-eGFP-PICV3739 at an MOI of 1 and sgRNA coverage of >500 in DMEM + 2% (v/v) FBS + 20 mM HEPES media. 24 hours post infection (hpi) cells were washed and media was changed and supplemented with 5 mM NH_4_Cl (Sigma-Aldrich, A9434) and 1% (v/v) penicillin-streptomycin (Gibco, 15140122). Cells were washed and treated with media supplemented with 5 mM NH_4_Cl every day for 11 days. Following NH_4_Cl treatment removal for 24 hr, half of cells were harvested, and cells were frozen as round 1 infection, and the other half were reseeded overnight for round 2 infections. Reseeded cells were infected at an MOI of 3. At 24 hpi cells were washed and media was changed with no NH_4_Cl. Surviving cells grew for an additional 24 days before harvesting all cells and storing at -80°C as a cell pellet for gDNA extraction and sequencing.

gDNA was harvested from reference and VSV-eGFP-PICV3739 selected cells using a Machery-Nagel Nucleospin Blood Kit (Takara, 740950). Cell pellets were treated with Proteinase K and lysed overnight at 70°C. Lysates were treated with 0.038 mg/ml RNaseA (Takara, 740505) for 5 min at room temperature, and gDNA was precipitated and purified according to manufacturer’s protocol (Takara, 740950), with the exception of samples being eluted with 70°C prewarmed elution buffer and incubated for 5 min at room temperature. Eluted gDNA was subjected to additional purification using a Zymo PCR inhibitor removal kit (Zymo Research D6030). gDNA was quantified using a Qubit fluorometer (Qubit 1X dsDNA HS Asssay Kit, Invitrogen, Q3231). Amplicon PCR and deep sequencing on a HiSeq (Illumina) were carried out at the Broad Institute (Genetic Perturbation Platform, Broad Institute of MIT and Harvard, Cambridge, MA). Enrichment of sgRNAs, as compared to unselected reference cells, was calculated using the MAGeCK software pipeline.^61,62^

### Plasma membrane targeted loss-of-function CRISPR-Cas9 Library Generation (surfaceome)

Four sgRNAs targeting 1,146 plasma membrane protein genes were identified from the genome-wid loss-of-function Brunello CRISPR library,^63^ along with 50 nontargeting control sgRNAs. lentiCRISPR-v2-PuroR containing sgRNAs were provided by the Genome Engineering and iPSV core (GEiC) at Washington University.^64^ Lentivirus was generated by Turbofectin 8.0 transfection of HEK293T cells supplemented with psPAX2 and pMD2.5 according to manufacturer’s protocol. Supernatants were harvested 2 days post-transfection, clarified by centrifugation and 0.45 µm filtration, and stored -80℃. 2.4x10^7^ HAP1 cells were transduced at an MOI of 0.3 with the lentivirus library and subsequently selected with puromycin (2 µg/mL) for 10 days.

### Plasma membrane-targeted loss-of-function CRISPR screening

Screening was done similarly as described for the whole-genome loss-of-function CRISPR screens. In brief, 4000 sgRNA coverage of the library was infected at an MOI of 3 with either VSV-PICV3739, VSV-PARV, or VSV-FLEXV for 1 hr in IMDM + 2% (v/v) FBS + 20mM HEPES, rocking every 10 min. Virus was then removed, and cells were supplemented with 15 mL of IMDM + 2% (v/v) FBS + 20 mM HEPES. At 24 hpi, cells were washed 1 time with media, and 5 mM NH_4_Cl was added to cells. Cells were washed daily, and new media supplemented with 5 mM NH_4_Cl was added for 9 more days. At 10 dpi, new media was added, and cells were grown without NH_4_Cl for 2 days before harvesting half of cells for gDNA and reseeding half to test for permissivity. The following day cells were infected with their corresponding virus at an MOI of 5. No further infection of the screened cells was noted, and no further rounds of infection were harvested or conducted. gDNA was harvested as described for whole-genome loss-of-function CRISPR screens. The sgRNAs from gDNA were amplified using a pooled forward primer (5’AATGATACGGCGACCACCGAGATCTACACCTGATGACACTCTTTCCCTACACGACGCTCTTCCGATCTN1-6TTGTGGAAAGGACGAAACACCG 3’) and a barcoded reverse primer (5’ CAAGCAGAAGACGCATACGAGATN8GTGACTGGAGTTCAGACGTGTGCTCTTCCGATCTCCAATTCCCACTCCTTTCAAGACCT 3’). The amplified cDNA was purified by gel extraction and sequenced by Illumina MiSeq by the Genome Access Technology Center at the McDonnel Genome Institute. sgRNA enrichment analysis done by the MAGeCK pipeline. ^61,62^

### Chimeric VSV infections for quantification

Cells were infected with indicated chimeric VSV at the stated MOI for 1 hr at 37°C, virus was then aspirated and media was changed and further incubated at 37°C for 5 hr. At 6 hpi, cells were trypsinized (Gibco, 1540054) and fixed in final concentration of 2-5% (v/v) formaldehyde (Sigma-Aldrich, 8187081000) for 15 min at room temperature, washed twice, and resuspended in 1X PBS. Cells were then subjected to flow cytometry analysis, collecting 10,000 singlets/condition and data was analyzed for cells, singlets, and %GFP+ (gating determined by <1% GFP+ in mock infected cells) using FlowJo (version 10) software.

### Soluble protein design, expression, and purification

The ectodomain of CD164, encompassing residues 1 through 162 was cloned as described previously,^27^ into the pcDNA3.1 vector (Addgene, 117272), generating a construct under a CMV promoter and with an N-terminal CD5 signal peptide in which sCD164 is fused to a C-terminal human IgG1 constant region (Fc) with a FactorXa linker (sCD164-Fc). Additionally, the ectodomain of hTfR1 (hTfR1-Fc) was designed and generated with domain boundaries as previously published.^65^ The CRD alone, encompassing residues 58 through 110, was cloned into the same vector as the sCD164 to generate the sCRD-Fc construct with the N-terminal CD5 signal peptide and C-terminal human Fc. sCD164 mutants were generated using Q5 site directed mutagenesis of each amino acid individually, resulting in sCRD-Fc. Clade A NWA GP1 constructs were cloned into a pTwist-CMV-ßGlobin mammalian expression vector (Twist Bioscience Inc.) under a CMV promoter with a N-terminal CD5 signal peptide with a C-terminal twin-strep tag fused to the GP1 with a Gly-Ser linker. GP1 truncations were designed based on multiple sequence alignment (Clustal Omega^66–68)^ and Alphafold3^48^ predicted structure alignment with the previously published MACV sGP1,^65^ residues as follows: PICV (77-264), PARV (77-266), FLEXV (77-266). The subsequent resulting proteins as denoted as: PICV sGP1-twinstrep, PARV sGP1-twinstrep, and FLEXV sGP1-twinstrep. Gibco’s Expi293 expression system was used to generate soluble proteins following manufacturer’s protocols (ThermoFisher Scientific, A14635). 18 hr after transfection, cells were enhanced per manufacturer’s protocol, and temperature was decreased to 31°C. At 5 days post-transfection, supernatants were harvested and clarified by centrifugation and filtration through a 0.45 μm filter. For Fc fusion proteins (sCD164, sCRD, hTfR1, or mutant sCD164), Protein A agarose resin (GoldBio, P-400-5) was added to clarified supernatants and gently agitated overnight at 4°C. Fc-conjugated proteins were purified by gravity filtration affinity chromatography (BioRad, 7321010), washed with 1 M Tris HCl at pH 8.0, and eluted with 0.2 M Glycine pH 2.2, and neutralized in 1:10 1 M Tris HCl pH 8.8. For twin-strep fusion proteins (sGP1s) Strep-Tactin XT 4Flow Resin (IBA Lifesciences, 2-5010-025) was stacked and washed with Buffer W (100 mM Tris-Cl, 150 mM NaCl, 1 mM EDTA pH 8.0) in a gravity filtration affinity chromatography column, filtered supernatants were then poured over the columns and washed in Buffer W and eluted with 50 mM biotin in Buffer W per manufacturers protocol. All purified proteins were dialyzed overnight in 1X PBS at 4°C in a dialysis cassette (10,000 MWCO, Slide-A-Lyzer 10K, Thermo Fisher Scientific, 66382), then concentrated with Amicon Ultra centrifugal filters (10,000 MWCO, MilliporeSigma, UFC901008). Proteins were stored at -80°C after flash freezing in liquid nitrogen. Protein quality was determined by 4-15% gradient SDS-PAGE gel (BioRad, 4561094) and staining with Coomassie stain (0.1% (w/v) Coomassie R350, 40% (v/v) Methanol, 10% (v/v) Acetic Acid). Prior to protein gel, some samples were treated with PNGaseF to remove *N*-linked glycosylations using manufacturer’s protocols (New England BioLabs, P0704) under denaturing and nondenaturing conditions to further ensure protein quality.

### BioLayer interferometry (BLI)

BLI experiments were performed on a GatorPlus BLI. Experiments were performed with universal buffer 2 composed of 20 mM Tris, 20 mM Bis-Tris, 20 mM HEPES, 500 mM NaCl, 1 mM CaCl_2_, 1 mM MgCl_2_, and 0.01% Tween-20 at indicated pH.^69^ For each binding assay, sCD164-Fc, hTfR1-Fc, sCRD-Fc, or mutant CD164, were first immobilized on GatorBio anti-human IgG Fc Gen II (HFCII) probes (GatorBio,160024) for 300 sec. Following capture of receptor-Fc, probes were baselined in buffer for 60 sec, followed by association with sGP1-twinstrep or antibody (for sCD164 association was done with mouse anti-human CD164 antibody N6B6 (BD Biosciences, 551296); for hTfR1 association was done with mouse anti-human CD71 (BD Pharmigen, 555534)) at indicated concentration for 180 sec then dissociation for 300 sec. All corresponding steps were conducted at indicated pH throughout the experiment.

Data were exported and pre-processed using a custom Python script. Data pre-processing included alignment of association and dissociation phases, reference subtraction, baseline adjustment, and spike correction. For determination of kinetic binding affinities, maximum response (R_max_), on-rate (k_on_) and off-rate (k_off_) were fit to either a 1:1 or 1:2 Langmuir site binding model represented by the integrated rate equation (BIAevaluation Software Version 3.0, Biacore AB). For determination of equilibrium binding affinities (Scatchard, Hill-Langmuir) and pH binding curves, pre-processed data were exported and analyzed and plotted in Prism. Python scripts used for pre-processing and analysis are available on GitHub (https://github.com/kaszubat/CD164-CANW-BLI).

### sGP1 cell binding and anti-CD164 antibody staining assays

CD164^KO^ or CD164^KO^+WT-CD164^PM^ cells were detached with 5 mM EDTA (ThermoFisher Scientific, J62948-A1) and blocked by washing three times with 1X PBS (Corning, MT-46013CM) + 7% (w/v) Bovine Serum Albumin (BSA, Sigma-Aldrich, A2153) at 4°C. Cells were then stained at indicated pH in 1X PBS + 1% (w/v) BSA for 30 min on ice with decreasing concentrations (5 μM to 0.019 μM) of PICV sGP1 labeled with AlexaFluor 647 carboxylic acid, succinimidyl ester (Invitrogen, A2006) following manufacturer’s protocols and clean-up with a Zeba spin desalting column (ThermoFisher Scientific, 89882). Alternatively, cells were stained with 1:1000 dilution of a polyclonal sheep anti-human CD164 antibody (“polyclonal antisera,” R&D systems, AF5790) at pH 7.4. Cells were then washed three times with 1X PBS + 1% (w/v) BSA at appropriate pH at 4°C. Cells stained with the anti-CD164 antibody were then stained with an AlexaFluor 647 donkey anti-sheep IgG H&L (Abcam, ab150179) for 30 min on ice and then washed three times with 1X PBS + 1% (w/v) BSA at pH 7.4 at 4°C. Cells were then maintained at respective pH on ice and subjected to flow cytometry analysis (Beckman Coulter Cytoflex S), collecting 50,000 singlets/condition and analyzed for cells, singlets, and AF647 mean fluorescent intensity (MFI) using FlowJo (version 10) software.

### Thermal stability assays

Stability of soluble PICV sGP1 and mutants were determined through thermal stability assays (Applied Biosystems, 4461146) following STAR protocols^70^ utilizing a ThermoFisher Scientifics QuantStudio 3. Protein was diluted to 5 μM in Phosphate Buffered Saline (PBS) with 1 mM CaCl_2_ and 1 mM MgCl_2_ and mixed with 8X dye and 5 μL of thermal shift buffer to a total volume of 20 uL. Derivative of fluorescence at temperature was determined through melt curve analysis on QuantStudio Design & Analysis 2.8.0 Software and plotted with Prism.

### Acid bypass and fusion assay

Indicated HeLa cells (WT, CD164^KO^ or CD164^KO^+WT-CD164^PM^) were pretreated with 5 nM of Bafilomycin A1 (BafA1, ThermoFisher Scientific, J61835-M) or DMSO for 1 hr on ice. Indicated virus was then added to cells at an MOI of 3 in the presence of 5 nM BafA1 or DMSO (Sigma-Aldrich, D8418) for an additional hour on ice. Virus was then removed, and fusion was forced at the plasma membrane using 37°C DMEM + 10 mM MES with 5 nM BafA1 or DMSO at the indicated pH for 15 min at 37°C. pH media was then removed and replaced with DMEM + 2% (v/v) FBS + 20mM HEPES with 5 nM BafA1 or DMSO and incubated at 37°C for 5 hr. Infections were fixed, processed, and analyzed as described in “chimeric VSV infections.”

### Immunostaining

Indicated cells were seeded on autoclaved 1.5 glass 12 mm round coverslips (Warner Instruments, 64-12R15) and incubated overnight. Before infection, cells were treated with cycloheximide (CHX, Sigma-Aldrich, C7698) for 30 min at 37°C at concentrations of 11.1 μM (for confocal microscopy) or 100 μM (for widefield microscopy). Cells were then infected with the indicated virus at an MOI of 400 in the presence of cycloheximide for 4 hr (confocal) or 2 hr (widefield) at 37°C. Following infection, cells were stained with 5.0 μg/mL of AlexaFluor 647 wheat germ agglutinin (Invitrogen, W32466) for 15 min at room temperature, washed with 1X PBS, and fixed with 4% (v/v) methanol-free paraformaldehyde (ThermoFisher Scientific, 28908). Cells were then permeabilized and blocked in 1X PBS supplemented with 3% (w/v) BSA and 0.1% (w/v) saponin (Sigma-Aldrich, SAE0073) for 30 min at room temperature. Cells were then stained with 1:1000 mouse anti-VSV-M antibody 23H12^46,47^ in wash buffer (1X PBS, 1% (w/v) BSA, 0.1% (w/v) saponin) for 1 hr at 37°C in a humidity chamber. Coverslips were washed three times in wash buffer, stained with AlexaFluor 594 goat anti-mouse antibody (Invitrogen, A-11032) and 600 nM DAPI (Invitrogen, D1306) for 15 min at 37°C, and mounted with ProLong Glass Antifade (Invitrogen, P36980) on glass slides (Electron Microscopy Sciences, 71883-05).

### Image acquisition and analysis

Confocal imaging was performed on a Zeiss Confocal LSM980 Airyscan inverted microscope. Images were acquired with a Plan-Apochromat 40×/1.4 water DIC objective lens with Z-stacks acquired for the entire depth of the cell at a step size of 0.18 µm. Colocalization analysis was performed on individual cells using the JACoP plugin in Fiji.^71,72^ Background fluorescence was removed via Otsu thresholding. Masked images were analyzed for Pearson correlation (measuring the degree of correlative variation in pixel intensity ranks) and Manders coefficient (measuring intensity co-occurrence).

Widefield microscopy was performed in a Zeiss Cell Observer Microscope spinning disk inverted microscope. Images were acquired as 4 × 4 tiles using a Plan-Apochromat 40x/1.4 water immersion objective and stitched using Zen Blue software. The resulting images contained approximately 400 cells each, a single representative tile is shown.

### CRD Site-Saturation-Variant Library Generation

Gene blocks containing the ectodomain of CD164 with all 52 amino acids independently mutated within the CRD to 20 other possible amino acids (Y, W, F, H, R, K, E, D, Q, N, T, C, S, P, M, V, I, L, G, A) was generated by Twist Bioscience Inc. Gene blocks were cloned via Gibson Assembly into the lentiviral cDNA expression plasmid pCW62-Puro (Harvard plasmid repository No. EvNO00438621) generating the plasmid library (pCW62-CRD-SVL-Puro). 30 colonies were picked and sequenced by Sanger Sequencing, with all sequences coming back with different individual amino acids mutated, demonstrating diversity in the plasmid library. DH10ß electrocompetent *E.coli* (New England BioLabs, C3019l) were electroporated with pCW62-CRD-SVL-Puro library with a final estimated coverage of 3.03x10^10^ and grown at 32°C for 18 hr. Plasmid was prepared with Machery-Nagel Xtra Midi kit for transfection-grade plasmid DNA (Machery-Nagel, 740410). Diversity of the library was confirmed by sequencing on an Illumina MiSeq instrument. Lentivirus was generated and HeLa CD164^KO^ cells were transduced as described under “generation of lentiviral addback cell lines,” at an MOI of 0.1 in an estimated coverage of 816. One day after spinoculation cells were combined and expanded, two days after spinoculation cells were treated and maintained in 2 μg/mL of puromycin.

### CRD site-saturation-variant screening

Prior to screening a reference library of >1,000 coverage was harvested and sequenced to confirm diversity of the library and for comparison of sorted screens.

To screen for amino acids that mutate the CRD and impact binding of PICV sGP1(GP1-) of the CRD, the Site Variant library was first Fluorescence-Activated Cell Sorted (FACS) (BD FACSSymphony S6, Department of Pathology and Immunology Flow Cytometry and Fluorescence Cell Sorting Core, Washington University School of Medicine) for plasma membrane expression of CD164 with two anti-CD164 monoclonal antibodies: AlexaFluor 594 labeled N6B6 (purified mouse anti-human CD164 antibody N6B6, BD Biosciences, 551296) and AlexaFluor 488 labeled 67D2 (purified mouse anti-human CD164 antibody 67D2, BioLegend, 324802). Cells were then expanded and maintained in 2 μg/mL of puromycin. Expanded CD164+ cells were sorted for lack of binding by AlexaFluor 647 labeled PICV sGP1 (AF647 negative population) at pH 6.0. To screen for CRD mutations that retain binding of PICV sGP1 (GP1+) the CRD-SVL was sorted for binding of AF647-PICV sGP1 at pH 6.0. To screen for CRD mutations that impact the binding of N6B6, for the identification of the N6B6 epitope, the CRD-SVL was sorted for binding of AlexaFluor 594 labeled N6B6 antibody and maintained as described. Genomic DNA of library and screens were then isolated using DNeasy Blood and Tissue Kits for DNA Isolation per manufacturer’s protocol (Qiagen, 69506).

First round Q5 PCR reactions were conducted on a total of 4 μg of individually isolated gDNA to amplify the CRD of CD164, 5’ end of the primers containing Illuminia sequences followed by 5 random positions (NNNNN). Resulting PCR product was gel extracted using Zymoclean gel DNA Recovery Kit (Zymo Research, D4001). The product then underwent second round indexing with Q5 PCR with TruSeq Indexing primers on 250 ng of gel extract PCR product. Resulting product was gel extracted and analyzed for quantity and purity by Qubit and Bioanalyzer respectively. Libraries were sequenced on an Illumina Miseq instrument and analyzed using the bcsubamp and diffsel tools of the dms_tools2 package.^73^ Differential selection values were plotted as heat maps using the ggplot2 package in R.^74^ Raw sequencing files have been deposited in the Sequence Read Archive (Accession PRJNA143394).

### CD164 addback cell line expression and quantification

CD164 expression in CRD mutant cell lines generated by lentivirus addback in CD164^KO^ HeLa cells was confirmed by western blotting and immunofluorescence staining. CD164 expression by immunofluorescence was done as described in “GP1 Cell Binding and anti-CD164 Antibody Staining Assays.” For western blots, cells were lysed with Pierce RIPA lysis and Extraction Buffer (ThermoFisher Scientific, 89900) following manufacturers protocol and then normalized for protein quantity with Peirce detergent compatible Bradford reagent (ThermoFisher Scientific, 23246). Samples were diluted in 4X Laemmli (BioRad, 1610747) + 50 mM Dithiothreitol (DTT, Invitrogen, D1532) and denatured by boiling for 10 min at 100°C. 4.7 μg of sample was run on a 4-15% gradient SDS-PAGE gel (BioRad, 4561094) and transferred to a nitrocellulose membrane. Membranes were then blocked with 1X Tris-Buffered Saline with 1% (v/v) Tween-20 (TBST) and 5% (w/v) milk for 1 hr at room temperature then stained overnight at 4°C with 1:1000 dilution of a polyclonal sheep anti-human CD164 antibody (R&D systems, AF5790) in TBST + 5% (w/v) milk. Membranes were then washed three times with TBST and stained with 1:1000 dilution of donkey anti-sheep IgG H&L HRP antibody (Abcam, 6900) in TBST + 5% (w/v) milk for 1 hr at room temperature and washed three times with TBST. Membranes were then exposed to Pierce Enhanced Chemiluminescence (ECL) Western Blotting Substrate (ThermoFisher Scientific, 32106) and imaged using a BioLegend ChemiDoc MP Imaging System. Quantification of western blotting was done by area under the curve (AUC) analysis with ImageJ, comparing CD164 AUC to actin AUC for respective sample then the relative expression of sample was compared to WT HeLa cells.

### ProteinMPNN analysis of PICV GP1

Structure-conserving designs of GP1 at the predicted GP1 and CRD interface (residues K213-C218 in GP1) were performed using ProteinMPNN.^75^ ProteinMPNN (v_48_020) was used to generate 50 sequences at a sampling temperature of 0.1 and 2.0. Highly divergent mutant sequences, perplexity score >2.5, were further screened using AlphaFold3 to down select designs for experiments that retained predicted ß-sheet structure, showing a representation of mutants generated. The new consensus sequence, RCTRSC, was generated by aligning all resulting sequences provided at the sampling temperature of 0.1.

### Statistical analysis

Statistical significance for microscopy of colocalization differences was determined by one-way analysis of variance (ANOVA) with nonparametric parameters (Kruskal-Wallis test) with Tukey’s multiple-comparison tests using Prism version 10 (GraphPad).

Additional statistical analyses were performed with Prism version 10 (GraphPad) include: one-way analysis of variance (ANOVA) with Tukey’s multiple-comparison tests, Area Under the Curve, and Two-tailed unpaired t-test, as described in respective figure legends. Significance indicated by asterisks (*) with: *\*p<*0.05, *\*\*p<*0.005, *\*\*\*p<*0.0005, *\*\*\*\*p<*0.00005.

