## Supplemental Figures for "CD164 is an endolysomal host factor for entry of Clade A New World Arenaviruses"

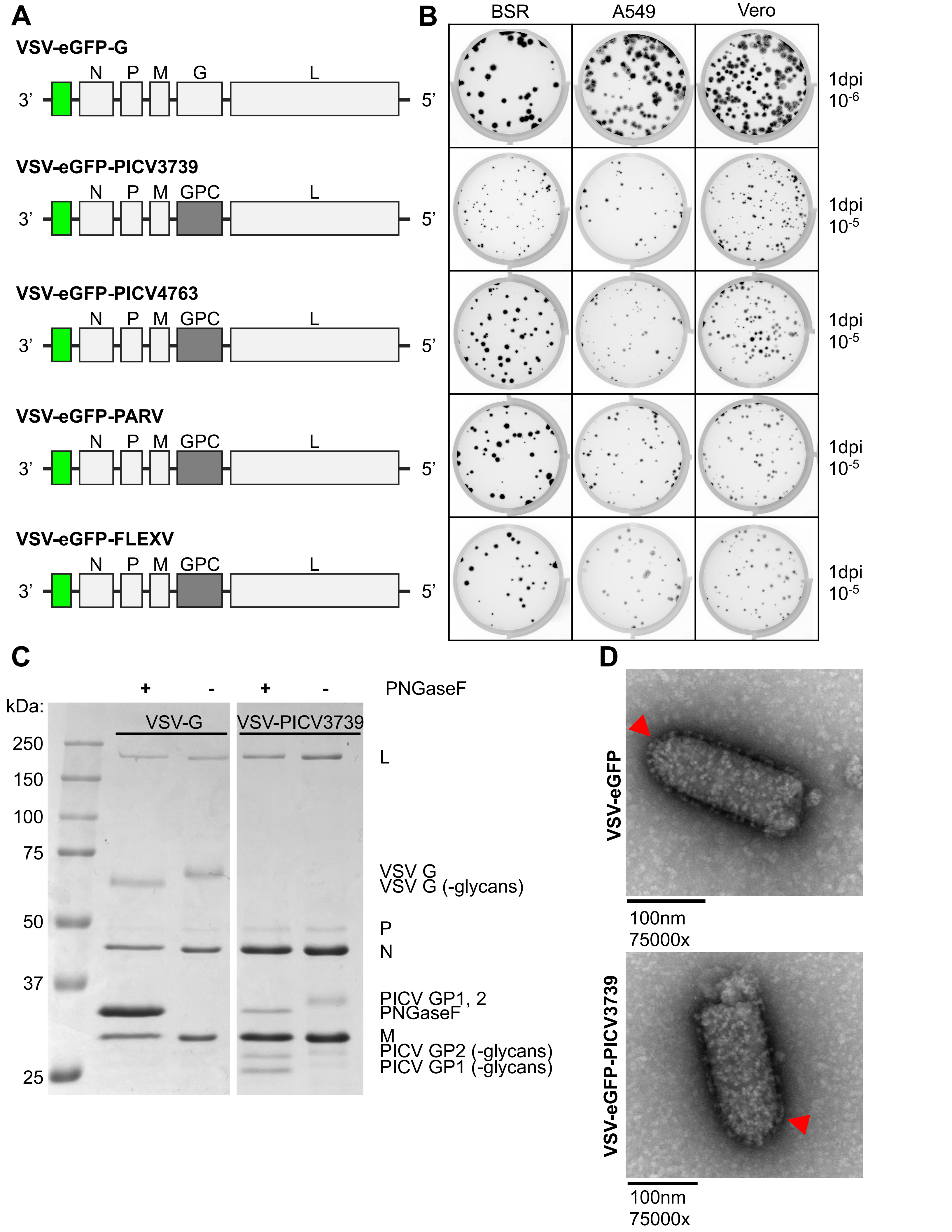
A
VSV-eGFP-G
N P M G L
3' 5'
VSV-eGFP-PICV3739
N P M GPC L
3' 5'
VSV-eGFP-PICV4763
N P M GPC L
3' 5'
VSV-eGFP-PARV
N P M GPC L
3' 5'
VSV-eGFP-FLEXV
N P M GPC L
3' 5'
B
BSR A549 Vero
1dpi 10-6
1dpi 10-5
1dpi 10-5
1dpi 10-5
1dpi 10-5
C
+ - + - PNGaseF
kDa:
VSV-G VSV-PICV3739
250
150
100
75
50
37
25
L
VSV G
VSV G (-glycans)
P
N
PICV GP1, 2
PNGaseF
M
PICV GP2 (-glycans)
PICV GP1 (-glycans)
D
VSV-eGFP
100nm
75000x
VSV-eGFP-PICV3739
100nm
75000x

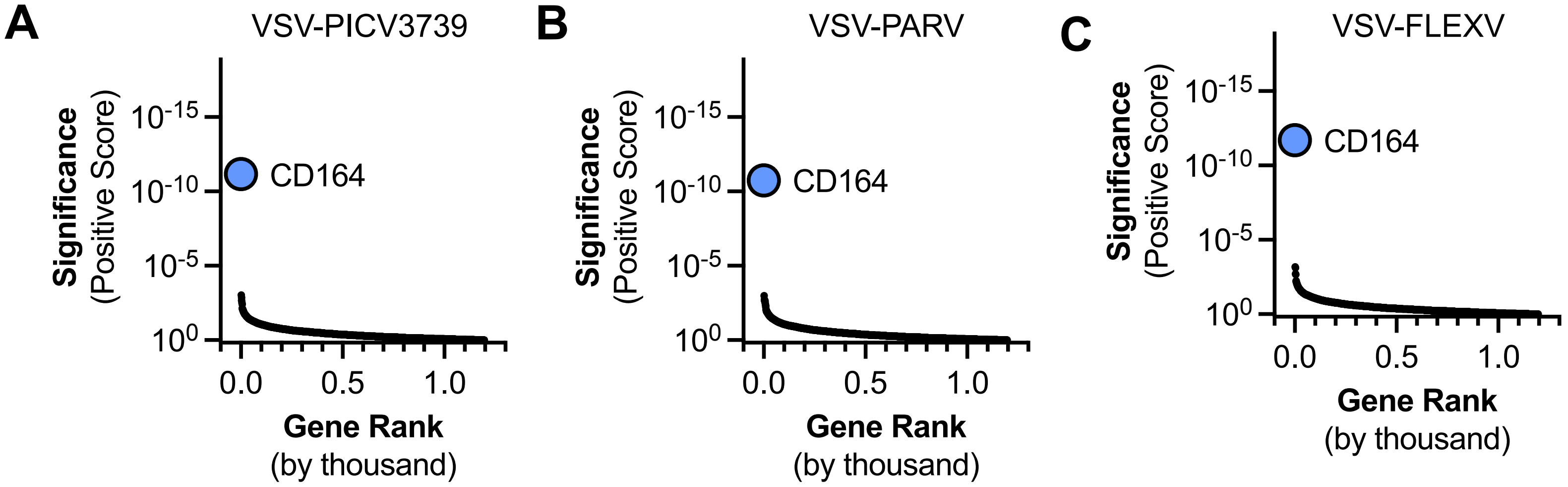

A
VSV-PICV3739
B
VSV-PARV
C
VSV-FLEXV
Significance
(Positive Score)
$10^{-15}$
$10^{-10}$
$10^{-5}$
$10^{0}$
CD164
0.0
0.5
1.0
Gene Rank
(by thousand)

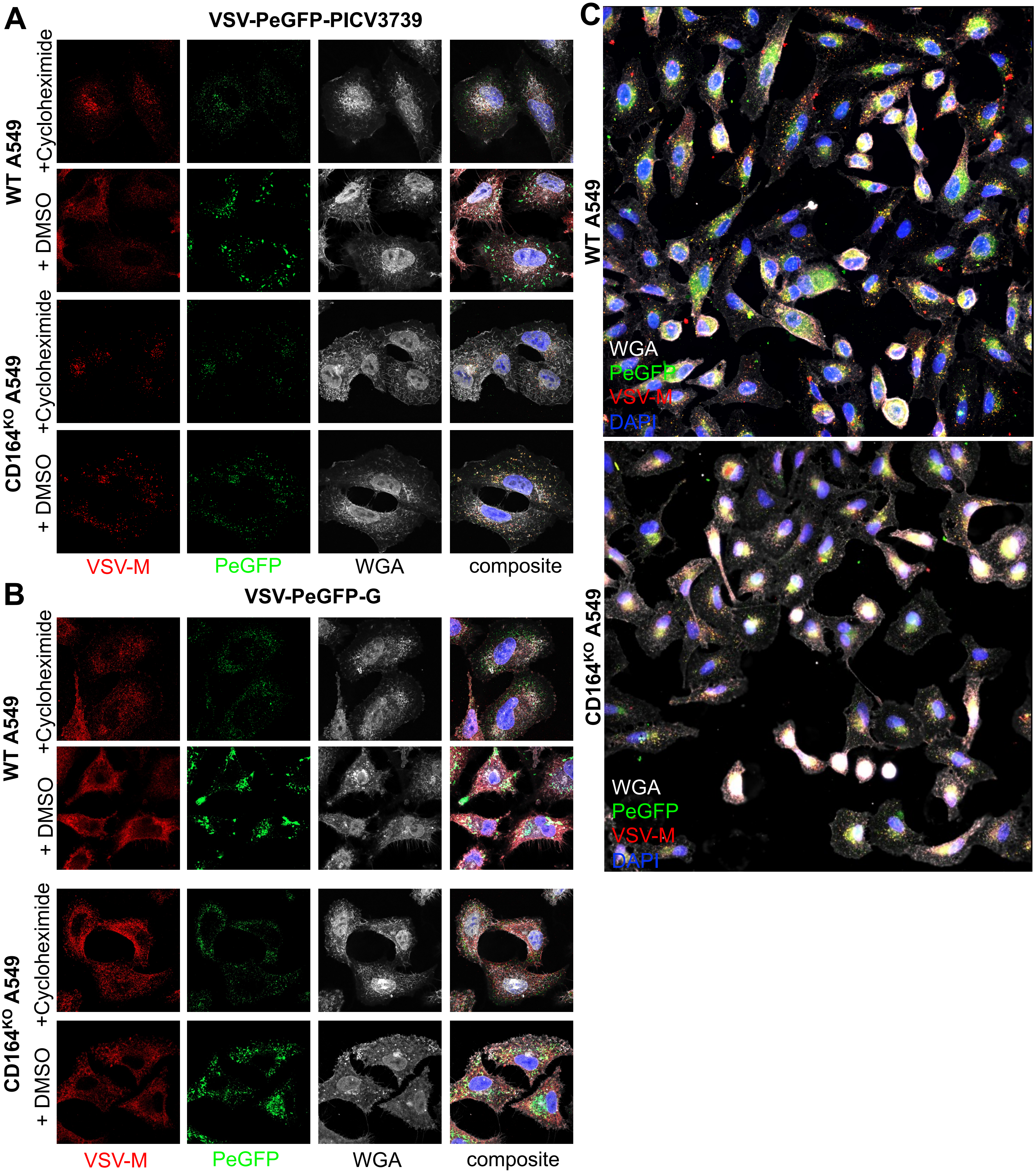
A
VSV-PeGFP-PICV3739
WT A549
+Cycloheximide
+ DMSO
CD164KO A549
+Cycloheximide
+ DMSO
VSV-M
PeGFP
WGA
composite
B
VSV-PeGFP-G
WT A549
+Cycloheximide
+ DMSO
CD164KO A549
+Cycloheximide
+ DMSO
VSV-M
PeGFP
WGA
composite
C
WT A549
WGA
PeGFP
VSV-M
DAPI
CD164KO A549
WGA
PeGFP
VSV-M
DAPI

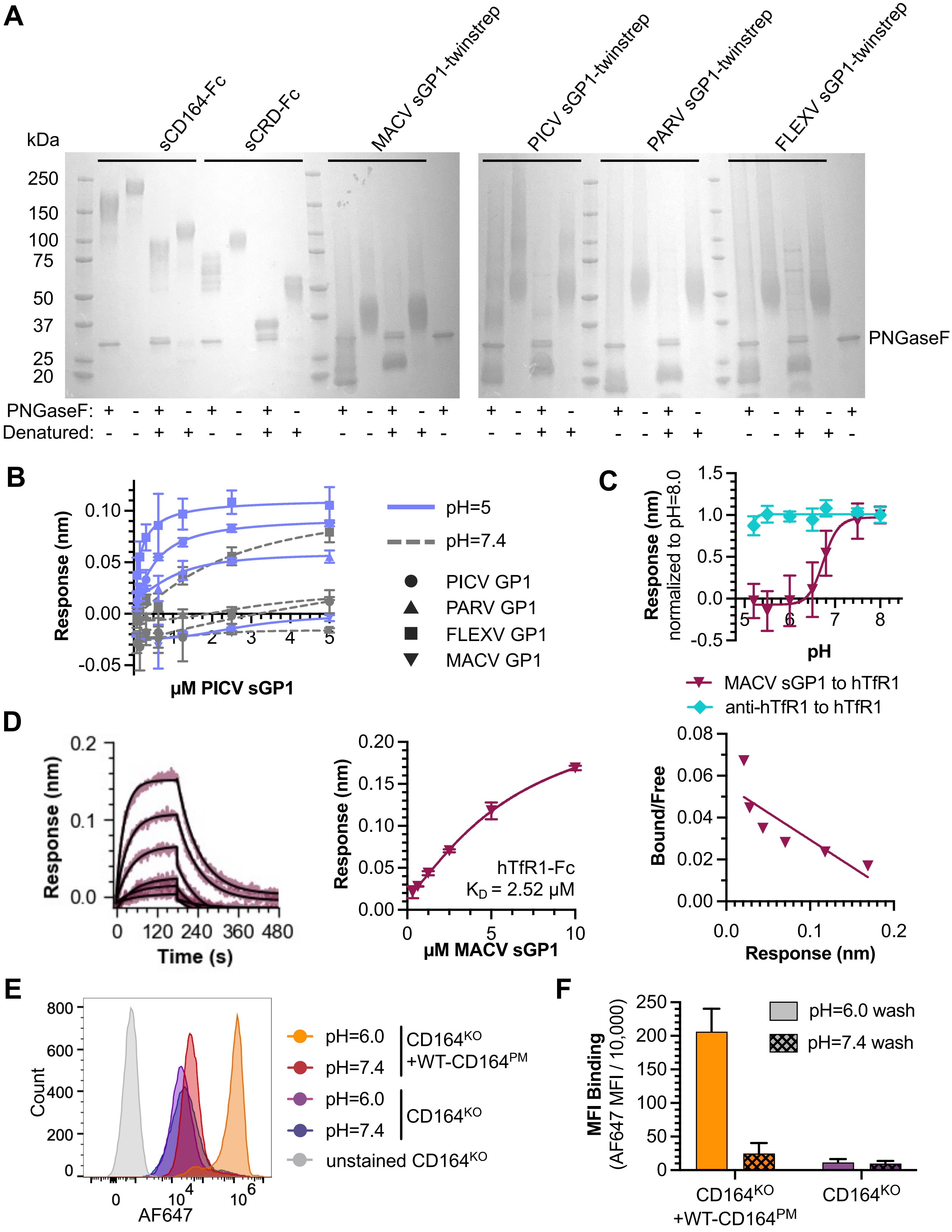

A
kDa
sCD164-Fc
sCRD-Fc
MACV sGP1-twinstrep
PICV sGP1-twinstrep
PARV sGP1-twinstrep
FLEXV sGP1-twinstrep
250
150
100
75
50
37
25
20
PNGaseF
PNGaseF:
Denatured:
B
Response (nm)
µM PICV sGP1
pH=5
pH=7.4
PICV GP1
PARV GP1
FLEXV GP1
MACV GP1
C
Response (nm) normalized to pH=8.0
pH
MACV sGP1 to hTfR1
anti-hTfR1 to hTfR1
D
Response (nm)
Time (s)
µM MACV sGP1
hTfR1-Fc
K$_D$ = 2.52 µM
Bound/Free
Response (nm)
E
Count
AF647
pH=6.0
pH=7.4
CD164$^{KO}$ +WT-CD164$^{PM}$
CD164$^{KO}$
unstained CD164$^{KO}$
F
MFI Binding (AF647 MFI / 10,000)
pH=6.0 wash
pH=7.4 wash
CD164$^{KO}$ +WT-CD164$^{PM}$
CD164$^{KO}$

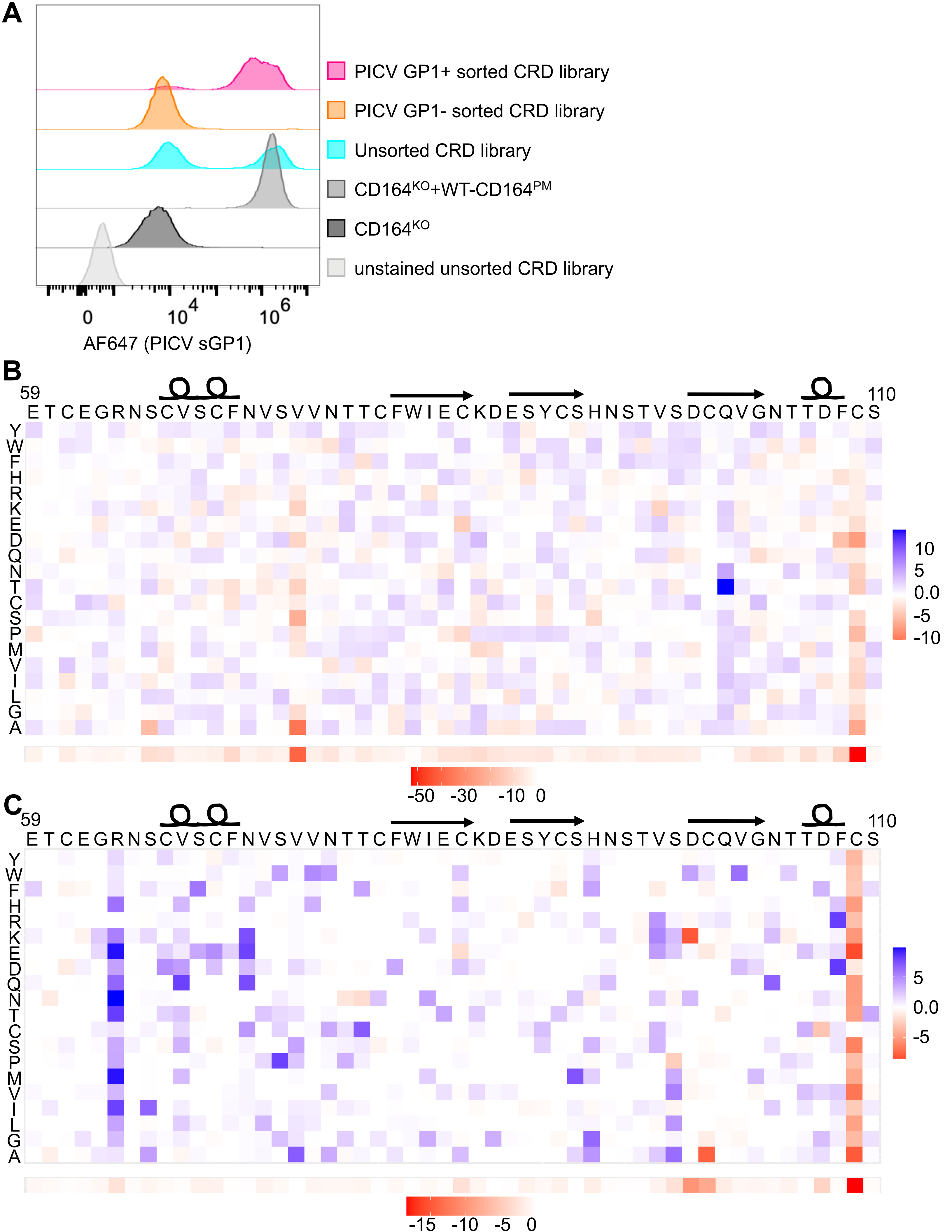
A
PICV GP1+ sorted CRD library
PICV GP1- sorted CRD library
Unsorted CRD library
CD164KO+WT-CD164PM
CD164KO
unstained unsorted CRD library
0
10^4
10^6
AF647 (PICV sGP1)
B
59
110
E T C E G R N S C V S C F N V S V V N T T C F W I E C K D E S Y C S H N S T V S D C Q V G N T T D F C S
Y W F H R K E D Q N T C S P M V I L G A
10
5
0.0
-5
-10
-50 -30 -10 0
C
59
110
E T C E G R N S C V S C F N V S V V N T T C F W I E C K D E S Y C S H N S T V S D C Q V G N T T D F C S
Y W F H R K E D Q N T C S P M V I L G A
5
0.0
-5
-15 -10 -5 0

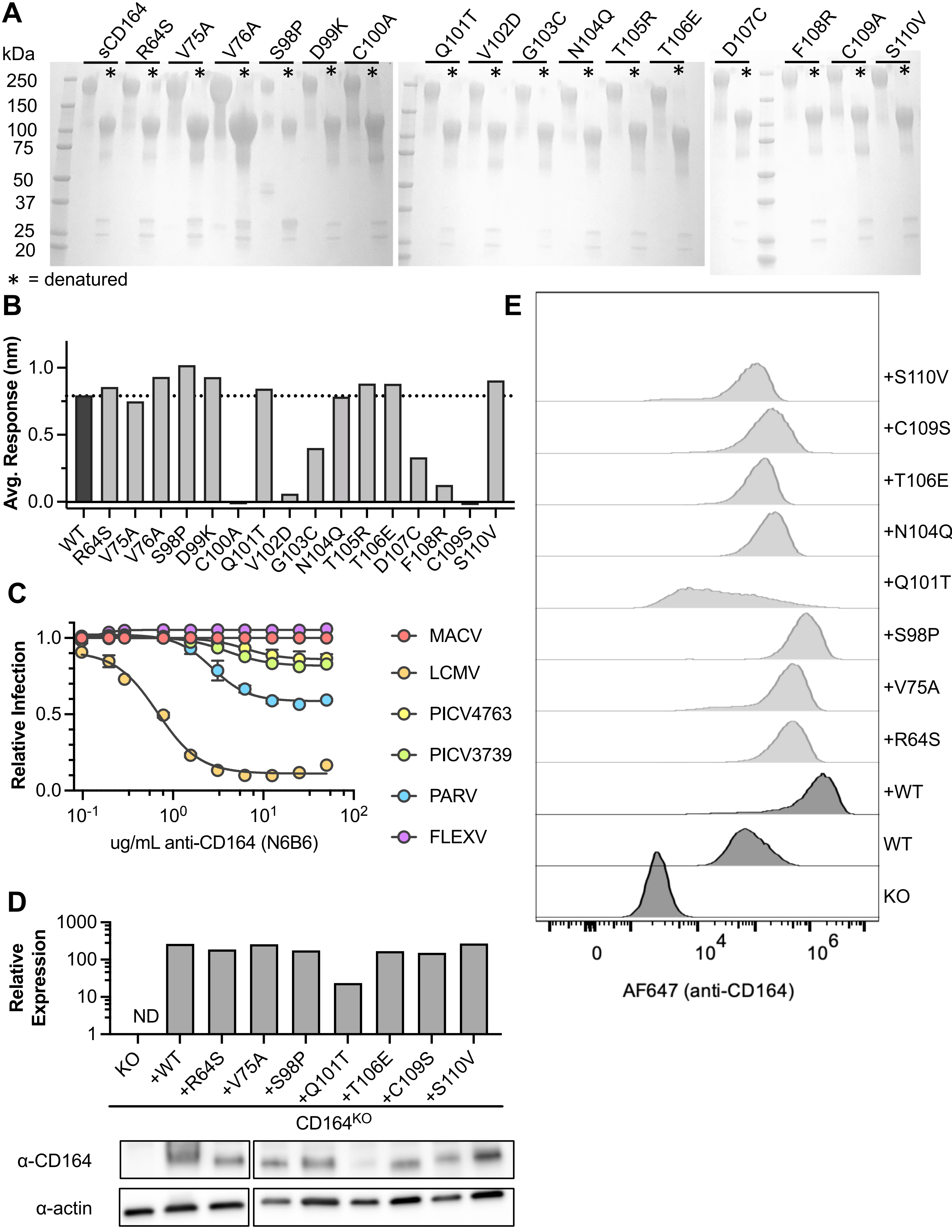
A
kDa
250
150
100
75
50
37
25
20
sCD164
R64S
V75A
V76A
S98P
D99K
C100A
Q101T
V102D
G103C
N104Q
T105R
T106E
D107C
F108R
C109A
S110V
* = denatured
B
Avg. Response (nm)
1.0
0.5
0.0
WT
R64S
V75A
V76A
S98P
D99K
C100A
Q101T
V102D
G103C
N104Q
T105R
T106E
D107C
F108R
C109S
S110V
C
Relative Infection
ug/mL anti-CD164 (N6B6)
MACV
LCMV
PICV4763
PICV3739
PARV
FLEXV
D
Relative Expression
1000
100
10
1
ND
KO
+WT
+R64S
+V75A
+S98P
+Q101T
+T106E
+C109S
+S110V
CD164KO
α-CD164
α-actin
E
+S110V
+C109S
+T106E
+N104Q
+Q101T
+S98P
+V75A
+R64S
+WT
WT
KO
AF647 (anti-CD164)

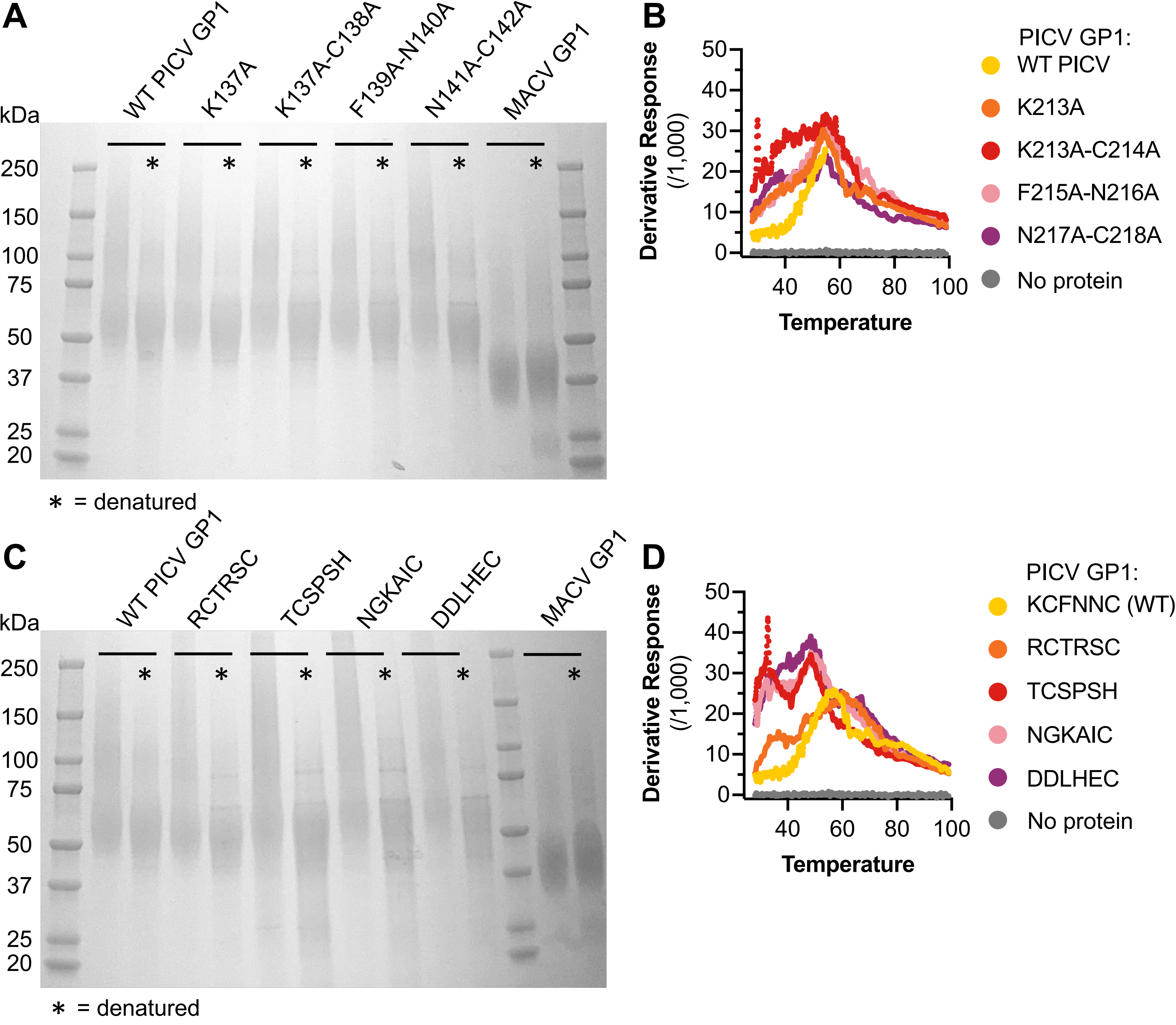


**E**

| | GPC | sequence | GPC |
|---|---|---|---|
| **MACV** | 202 | --GFHDPCE | 208 |
| **PICV3739** | 211 | VGKCFNNCS | 219 |
| **PICV4763** | 211 | VGKCFNNCS | 219 |
| **PARV** | 209 | VGKCFNNCS | 217 |
| **FLEXV** | 209 | VGKCDSRCS | 217 |
| **LCMV** | 214 | DGK.TTWCS | 221 |
